# High Throughput Meta-analysis of Antimicrobial Peptides for Characterizing Class Specific Therapeutic Candidates: An *in-silico* Approach

**DOI:** 10.1101/2024.09.15.613163

**Authors:** Anwesh Pandey, Raji Rajesh Lenin, Sumeet Patiyal, Piyush Agrawal

**Affiliations:** Department of Physics, Babasaheb Bhimrao Ambedkar University, Lucknow, India; Division of Medical Research, SRM Medical College Hospital & Research Centre, SRMIST, Kattankulathur, Chennai, India; Cancer Data Science Lab, National Cancer Institute, NIH, USA

## Abstract

The increasing incidence of antimicrobial resistance (AMR) is becoming a serious concern worldwide and requires newer drugs. Recent evidence has shown growing interest in developing peptide-based therapeutics to tackle AMR. In the current study, we performed a meta-analysis of nearly 8.6 million predicted antimicrobial peptides (AMPs) and assessed their antibacterial, antiviral, and antifungal activity. We created high-quality, class-specific datasets and performed several analyses, including amino acid composition, motif preference, physiochemical properties, etc. We observed significant differences in the residue composition, charge, molecular weight, pI value, and instability index of peptides among 3 AMP classes. We further developed multiple machine learning models to predict peptide activity using the dataset. Our Extratree model developed using composition-based features achieved the highest AUROC of 0.98 for antibacterial peptide (ABP), 0.99 for antiviral peptide (AVP), and 0.99 for antifungal peptide (AFP) prediction when tested on an independent dataset. Subsequent filtering of peptides based on moonlighting properties (toxicity, allergenicity, cell-penetrating ability, half-life, and secondary structure) yielded a list of peptides that exhibit substantial therapeutic potential. We further selected the top 10 peptides in each category based on their half-lives, predicted their 3D structures using ColabFold, a feature built into ChimeraX1.8 software, and used HDock to perform molecular docking analysis with a pathogenic protein selected from an organism in each class. Docking studies demonstrated strong interaction between peptides and proteins, and based on free energy, we ranked the peptides. Overall, we have put forth class-specific peptides with high therapeutic potential based on rigorous meta-analysis encompassing ∼8.6 million AMPs.

## Introduction

Diminutive and biologically active compounds produced by many organisms including animals, plants, humans and microorganisms during their innate immune responses are called antimicrobial peptides (AMPs) [1,2]. Infections or inflammations trigger the production of these peptides which are known for having different structure types and short sequences. One common feature of AMPs is that they have a broad range of activities that include pore/channel creation on microbial membranes to cause lysis [3,4]. It often involves interaction with lipid components present in microbial membrane leading to its destabilization and permeability changes [5]. AMPs have been extensively studied due to their effectiveness as traditional antibiotics that risk resistance escalation in the treatment of bacterial infections, viral diseases and cancer [6,7].

The rise in antibiotic resistance has made it necessary to find an alternative for conventional antibiotics – the use of antimicrobial peptide-based therapies is becoming more and more attractive. Major AMP-based drugs encompass LL-37, a human peptide exhibiting broad spectrum activity and immunomodulatory properties; Daptomycin, a cyclic lipopeptide interfering with bacterial membrane function; Nisin, a lantibiotic inhibiting cell wall synthesis as well as creating membrane pores; and Vancomycin-related peptides against bacterial cell wall precursors that are involved in AMP production [8]. These AMPs have wide activity range, multiple action mechanisms, and shorter development duration of resistance. Nevertheless, there remain some unanswered questions concerning potential toxicity, high costs of their manufacture and need for further research. To be able to effectively handle antimicrobial resistance better attempts are being made to increase their efficiency levels while at the same time minimizing any adverse effects that they may have on patients who use them by incorporating them into therapeutic regimens.

Antimicrobial peptides (AMPs) are categorized based on the pathogens they target and the way they work. They include antibacterial, antiviral, antifungal and antiparasitic peptides. One common aspect of many AMPs is their ability to destroy cellular membranes, although each subgroup has its own mode of action. For instance, defensins and nisin exemplify antibacterial peptides that often promote pore formation in bacterial membranes or inhibit cell wall synthesis while, enfuvirtide and Epi-1 demonstrate how antiviral peptides slow down viral entry interfering with viral envelopes [9]. Meanwhile, histatins and plectasin serve as examples of antifungal peptides that focus on fungal membranes or cell wall synthesis. In a similar way, magainin’s and defensins are instances where antiparasitic peptides target parasite membranes or metabolic pathways respectively [10]. Irrespective of their different targets, AMPs generally have broad-spectrum activities that are important for combatting multi-resistance challenges posed by microbes.

One common characteristic of AMPs is that they have membrane disruption as their primary mode of action, while the specific targets and actions may differ. In most cases, antibacterial, antifungal and antiparasitic peptides involve membrane damage whereas antiviral peptides often target either viral replication or entry. Every subclass of AMP has been adapted to fit pathogen structures or processes thus enabling them to bypass traditional resistance mechanisms hence their potential in developing new therapies against a wide range of microbial challenges [11].

Recent progress in the field of machine learning (ML) has had a profound impact on the analysis and advancement of antimicrobial peptides (AMPs), enriching their design, forecasting, and comprehension of underlying mechanisms. Forecasting the activity of AMPs has become more sophisticated with the utilization of ML models, such as support vector machines (SVMs), random forests, and deep neural networks (DNNs), for categorizing peptides based on established datasets to predict their antimicrobial efficacy [12]. Regression models are employed to anticipate the potency of peptides against specific pathogens by scrutinizing the correlation between peptide sequences and their biological functions. Deep learning methodologies, like convolutional neural networks (CNNs) and recurrent neural networks (RNNs), are adept at recognizing intricate patterns in peptide sequences, thereby enhancing prediction precision. The amalgamation of various models through ensemble techniques further amplifies performance in forecasting peptide activity [13].

The creation and improvement of AMPs have been revolutionized through generative models such as Generative Adversarial Networks (GANs), enabling the generation of novel peptide sequences possessing desired antimicrobial attributes by learning from existing datasets. Reinforcement learning aids in the iterative refinement of peptide sequences, assessing their effectiveness and optimizing their attributes. Automated design procedures leverage ML to streamline peptide synthesis, thereby minimizing the time and expenses associated with experimental endeavors and expediting the production of efficacious AMPs [14].

Insight into the mechanisms of action of AMPs is facilitated by ML techniques that extract and scrutinize features from peptide sequences and their interactions with microbial membranes, providing valuable information on how specific characteristics impact antimicrobial activities. Dimensionality reduction techniques like Principal Component Analysis (PCA) and t-Distributed Stochastic Neighbor Embedding (t-SNE) aid in visualizing and interpreting complex datasets to identify pivotal attributes driving peptide activity. The fusion of ML with molecular dynamics simulations offers an intricate understanding of peptide-membrane interactions and mechanisms of action within biological contexts [15].

ML classifiers are adept at predicting the toxicity and potential adverse effects of AMPs based on their sequences and structural properties. This proactive evaluation assists in identifying potentially harmful peptides before extensive experimental evaluations. Integrated methodologies that combine ML with cheminformatics and bioinformatics tools enhance the precision of toxicity forecasts, ultimately refining the safety profile of AMPs [16].

Incorporating ML techniques to fuse genomics, proteomics, and transcriptomics data provides a holistic perspective on peptide activity and interactions. This multi-omics strategy aids in comprehending how AMPs impact both microbial and host systems. Cross-validation with empirical data enhances the accuracy of ML forecasts and bolsters the reliability of AMP analysis. In conclusion, the strides made in ML are significantly propelling the realm of antimicrobial peptides by elevating activity prognostications, streamlining design processes, unraveling mechanisms, anticipating toxicity, and amalgamating diverse data reservoirs. These advancements are paving the way for pioneering peptide therapeutics and combating the escalating hurdle of antimicrobial resistance.

## Methodology

### 1. Dataset Creation

The dataset was downloaded from the recently published database AMPSphere by Santos-Junior et al. [17]. We downloaded data for nearly 8.7 million peptides as proposed by the authors in their original study and filtered them based on the following criteria: (i) high-quality only, (ii) with matched experimental data only, and (iii) peptides up to a length of 50 amino acids. This filtering resulted in 15,711 unique high-quality peptides, which we termed the “HQ_AMP_DS” dataset.

The activity of the peptides from “HQ_AMP_DS” was predicted for three major classes of AMPs: (i) antibacterial, (ii) antiviral, and (iii) antifungal. Activity was predicted using previously published machine or deep learning-based software. For antibacterial activity, we selected AntiBP3 [18], DRAMP 3.0 [19], PTPAMP [20], and iAMPpred [21]; for antifungal activity, we used AntiFP [22], PTPAMP [20], and iAMPpred [21]; and for antiviral activity prediction, we used AI4AVP [23], PTPAMP [20], and iAMPpred [21]. Software that was either not operational or very slow in providing output were not considered in this study.

As most of the tools require peptide sequences as input in ‘fasta’ file format, we converted the sequences using in-house scripts. We either pasted or uploaded these sequences as files to each webserver, selecting the default model, as recommended by the tools in their original paper, to predict the activity of a peptide. We calculated the tool’s performance in terms of sensitivity (accurately predicting positive classes) using Equation 1, based on the activity prediction results

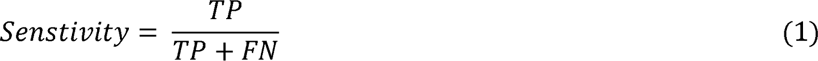

Where TP is True Positive and FN is False Negative.

Next, the peptides were categorized into their respective classes: antibacterial, antifungal, and antiviral. This classification was based on their predicted activity (positive or negative) with a cutoff value. For example, a peptide was considered to have antibacterial properties if it was predicted as positive by all four software tools and the cutoff value was above 0.8 (except for PTPAMP, where the cutoff used was 0.5, as most software values range between 0-1). Similarly, peptides were categorized into antiviral and antifungal classes. As many peptides might be common among different classes, we further filtered them to identify peptides unique to a particular class. These peptides were termed the positive dataset.

For the negative dataset, we followed three different approaches, which are discussed below:

Approach 1: Peptides were classified as negative, if they were predicted as non-antibacterial, non-antifungal, or non-antiviral based on a cutoff value. In case of non-ABPs cutoff value was <=0.2 except PTPAMP, with cutoff of -0.5 or less. For non-AFPs, cutoff was -1 for AntiFP, -2 for PTPAMP and 0.2 for iAMPPred. Likewise, for non-AVPs, cutoff was -0.1 for AI4AVP, -2 for PTPAMP and 0.2 for iAMPPred. The variation in cutoff selection was due to variability in the score provided by individual software. The peptides were further filtered out if they were common to any other two classes to obtain class-specific unique peptides.

Approach 2: In this approach, negative peptides were taken from published literature where they were used to develop in silico prediction tools. For example, the negative dataset used in developing antibacterial peptide prediction tools was taken as the negative dataset in the antibacterial section. After collecting all the data, only unique ones were kept.

Approach 3: Here, positive peptides of the other two classes were considered as the negative class for the third class. For example, peptides predicted to have antiviral and antifungal properties were considered the negative set for antibacterial peptides.

In total, we created 3 different datasets, where the positive peptides were common and only the negative set was different. We termed them High-Quality Dataset 1 (HQ_DS1) with positive peptides and negative peptides obtained from Approach 1; High-Quality Dataset 2 (HQ_DS2) with positive peptides and negative peptides obtained from Approach 2; and High-Quality Dataset 3 (HQ_DS3) with positive peptides and negative peptides obtained from Approach 3.

### 2. Analysis of Class Specific Peptide Properties

The following analyses were performed to understand the class-specific properties of the peptides.

#### 2.1. Amino Acid Composition Analysis

Amino acid composition represents the amino acid residue frequency in a given peptide/protein and plays an important role in elucidating its mechanism of action. Multiple studies have utilized this as a feature to develop machine-learning-based tools [24,25]. We computed the amino acid composition of positive and negative peptides for all three datasets associated with each class of AMPs using the following equation:

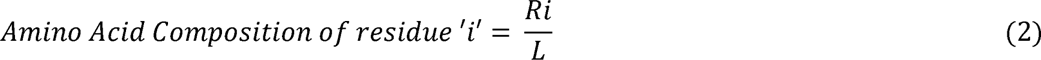

Where R_i_ represents number of residues of type ‘i’ and L represents the length of the sequence.

#### 2.2. Physiochemical Properties Analysis

We got the different physiochemical properties (mass, charge, pI value, and instability index) for each peptide from the AMPSphere metadata so that we could compare each property between the three predicted classes, which are ABP, AVP, and AFP. We presented each property individually in the form of a boxplot and used the Mann-Whitney test to determine the statistical significance of the distribution.

#### 2.3. Two Sample Logo Analysis

Using the Two Sample Logo (TSL) online tool [26], we generated sequence logos for each of the three datasets in each category. The sequence logo represents the amino acid residue with the x-axis, displays the bit-score for enriched residues on the positive y-axis, and displays the bit-score for depleted residues on the negative y-axis. This arrangement demonstrates the relative importance of each residue in each position. We chose the first ten residues of each peptide from the positive and negative classes since TSL required sequences in a fixed-length vector format. We conducted the analysis for both the N and C termini of the peptide.

#### 2.4. Motif Analysis

We used the MERCI software [27] to conduct motif analysis and characterize the patterns and motifs present in both the positive and negative classes of peptides. When running the software, we used its default settings. The tool compares the two sets of peptides (positive and negative) to characterize exclusive motifs in one set relative to the other. For instance, the tool considers ABPs as a positive set and non-ABPs as a negative set in order to obtain exclusive motifs in ABPs. In the input file order, however, we treat ABPs as negative classes and non-ABPs as positive classes, which enables us to extract exclusive motifs from non-ABPs. We analyzed the three datasets created for each class.

### 3. Machine Learning Analysis

#### 3.1 Feature Generation

It is a key step to transform amino acid sequences into numerical vectors for developing machine learning models. We used the Pfeature tool’s [28] composition module to compute several types of features. This includes features based on composition, such as the composition of amino acids, dipeptides, tripeptides, atoms, and diatoms; features based on physiochemical properties; and features based on Shannon entropy, such as the Shannon entropy of whole peptides (SEP), each residue (SER), and physical properties (SPR). We also calculated a number of other features, including distance distribution of residues (DDR), which shows how residues are arranged in space; residue repeat information (RRI), which looks at patterns of repetition within the sequence; pseudo-amino acid composition (PAAC) and amphiphilic pseudo-amino acid composition (APAAC), which are more advanced representations that take sequence and physicochemical properties into account; conjoint triad calculation (CTC), which focuses on the three residue groups; composition enhanced-transition distribution (CETD); and sequence order coupling number (SOC), which shows how transitions and couplings happen within the sequence; and Quasi-sequence order (QSO) feature capturing the subtle patterns present in the sequence. All these features collectively provide a comprehensive amino acid sequence representation, enabling robust training and evaluation of machine learning models.

#### 3.2 Model Development

In the present study, we developed and rigorously evaluated distinct machine learning models to classify peptides into 3 distinct classes of AMPs, i.e., ABPs and non-ABPs, AFPs and non-AFPs, and AVPs and non-AVPs, a crucial task for advancing peptide-based therapeutic research. We implemented seven distinct ML classifiers: (i) decision tree (DT), (ii) random forest (RF), (iii) logistic regression (LR), (iv) extreme gradient boosting (XGB), (v) Gaussian naïve Bayes (GNB), (vi) extra trees (ET), and (vii) support vector classifier (SVC) to explore the unique strength each classifier possesses. DT is well known for interpretability; RF and ET enhanced accuracy and robustness through ensemble methods, whereas LR offered insights into feature influence. XGB is known for optimized performance through boosting; GNB made simplifying assumptions that sometimes closely matched the data characteristics; and SVC excelled in handling high-dimensional data with effective decision boundaries. We implemented a widely used Python-based scikit-learn library [29] to develop models that were not only accurate but also robust and generalizable across different peptide classifications.

#### 3.3 Five-fold Cross Validation

We implemented stratified five-fold cross-validation to ensure the model’s robustness and accuracy. In this technique, data is divided into 5 equal sets, maintaining the percentage of classes, and four parts are used for training and the fifth one is used for testing. The process is repeated five times, where each set is used as a testing set once. In the end, average performance is computed post-5 iterations and provides a reliable estimate of model performance. The process reduces the risk of model overfitting and ensures that the model is generalizable to unseen data.

#### 3.4 Performance Evaluation Metrics

Model performance was evaluated using threshold-dependent and independent matrices. In the case of threshold-dependent, we computed sensitivity, specificity, accuracy, F1 score, Kappa, and Matthew’s correlation coefficient. Equations for these are provided in Equations 1 & 3-7. For threshold independence, we computed two widely used matrices, i.e., Area Under the Receiver Operating Characteristic Curve (AUROC) and the Area Under the Precision-Recall Curve (AUPRC). Complete details of these parameters are provided in the previous study [30,31]. Both the metrics were used to achieve a comprehensive model evaluation, ensuring they handle class imbalances effectively and make reliable predictions.

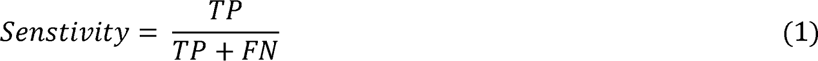

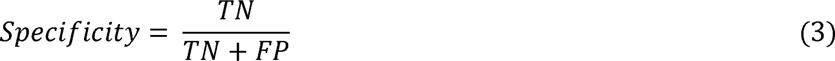

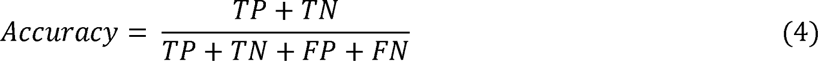

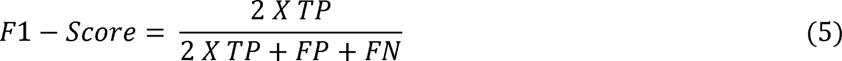

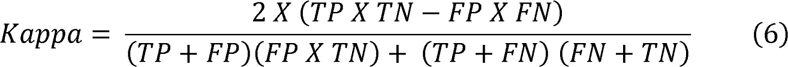

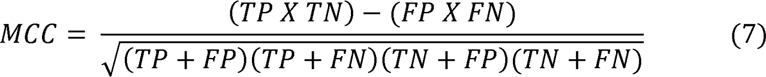

Where, TP stands for True Positive, TN stands for True Negative, FP stands for False Positive, and FN stands for False Negative.

### 4. Characterizing peptides with therapeutic potential

We performed multiple experiments to reduce the list of peptides acquired initially to get the high-quality candidates fulfilling various clinical properties for being a safer and more effective therapeutic candidate. Knowledge acquired over the years suggests that peptides used for therapeutic purposes exhibit a range of essential features, which include their ability to penetrate the pathogen’s cell membrane; should be non-toxic and non-allergic to the host; should have structural stability under physiological conditions; and an adequate half-life in the blood to ensure prolonged therapeutic action. The above-mentioned criteria are important to ensure that the peptide-based drugs can function effectively without causing adverse effects, making them suitable candidates for clinical applications.

We predicted and evaluated these traits using the recently developed tools, based on several biologically important features and machine learning approaches, each of which was specifically developed to assess the different properties of the peptides. For predicting allergenic properties, we used the AlgPred 2.0 online server [32]; to predict whether a peptide is toxic, we used the ToxinPred 3.0 server [33], and the CellPPD server [34] was employed to predict the cell-penetrating ability of the peptides. The ‘PLifePred’ webserver [35] was used to measure the half-life of peptides in the blood. We selected default parameters and models to perform these analyses. For predicting the enriched secondary structure in the peptides, we used the GOR4 online server [36]. This server takes peptide sequences as input and provides the content (%) of α-helix, β-sheet, random coil, etc., as output.

By analyzing these above-motioned key features, we systematically narrowed down the pool of peptides to characterize the peptides with elevated therapeutic potential. These peptides can be potential candidates for treating various diseases caused by bacteria, fungi, or viruses. The meticulous selection procedure and application of cutting-edge computational methods greatly increased the accuracy of our predictions, paving the way for possible therapeutic deployment and subsequent experimental confirmation.

### 5. Structure Prediction and Molecular Docking Analysis

The anti-bacterial, anti-fungal, and anti-viral peptide sequences were modelled using machine learning tools for a reliable structure. We used ColabFold [37] embedded in ChimeraX1.8 software [38] for predicting the structure of the peptides. For each peptide sequence, 5 models were generated, and the best model, based on Multiple Sequence Alignment (MSA), and predicted local distance difference test (pLDDT) values was chosen for further studies.

Next, as a case study for the docking experiments, we selected one protein from each class of AMPs. In case of bacteria, we selected the Streptococcus pneumoniae surface protein [PDB ID: 3ZPP]; for fungus, we selected Candida albican’s Als3 adhesin protein [PDB ID: 4LE8] and in case of virus, we selected SARS-CoV 2 main protease protein [PDB ID: 6LU7]. 3D-structures of these proteins were downloaded from the RCSB-Protein Data Bank [PDB] [39] and were prepared for the docking. We carried out docking calculations for each of the modelled peptides in each class with the corresponding protein and generated ten poses for each. HDock webserver [40] was employed to dock the peptides to their respective proteins. The pose with highest docking score was selected for the interaction analysis. While performing the docking calculations the native charges of the residues of each of the proteins was preserved, in lieu of the interactions being predominantly non-covalent in nature. The docked poses were visualised using the PyMOL software [41], and the information about the interacting residues was obtained using the PDBsum software [42], wherein chain-A represents the protein sequence, and chain-B represents the peptide sequence, respectively.

## Results

### Predicted AMPs possess multi-class functional properties

The authors of the original study suggest the predicted peptides as possible AMPs. However, AMPs can be roughly categorized into three classes: (i) antiviral (AVPs), (ii) antifungal (AFPs), and (iii) antibacterial (ABPs). Using previously created in silico methods for each class, we predicted class-specific peptide activity within the “HQ_AMP_DS”. We utilized iAMPpred, DRAMP, PTPAMP, and AntiBP3 to test for antibacterial properties. Results show that iAMPpred scored the highest, 69.19%, followed by AntiBP3 (60.87%), DRAMP (52.32%), and PTPAMP (12.04%). We employed AntiFP, PTPAMP, and iAMPpred to test for antifungal characteristics. As shown in Table 1, iAMPpred again demonstrated the best sensitivity at 58.52%, followed by PTPAMP at 53.74% and AntiFP at 20.43%. PTPAMP, iAMPpred, and AI4AVP were utilized to predict antiviral properties. In this case, iAMPpred demonstrated the maximum sensitivity (64.44%), trailed by AI4AMP (31.37%) and PTPAMP (21.82%). Performance of various software in classification of a peptide in positive or negative class is provided in Table 1. In general, most of the methods didn’t predict the class of peptides correctly with high degree of sensitivity.

**Table 1.**
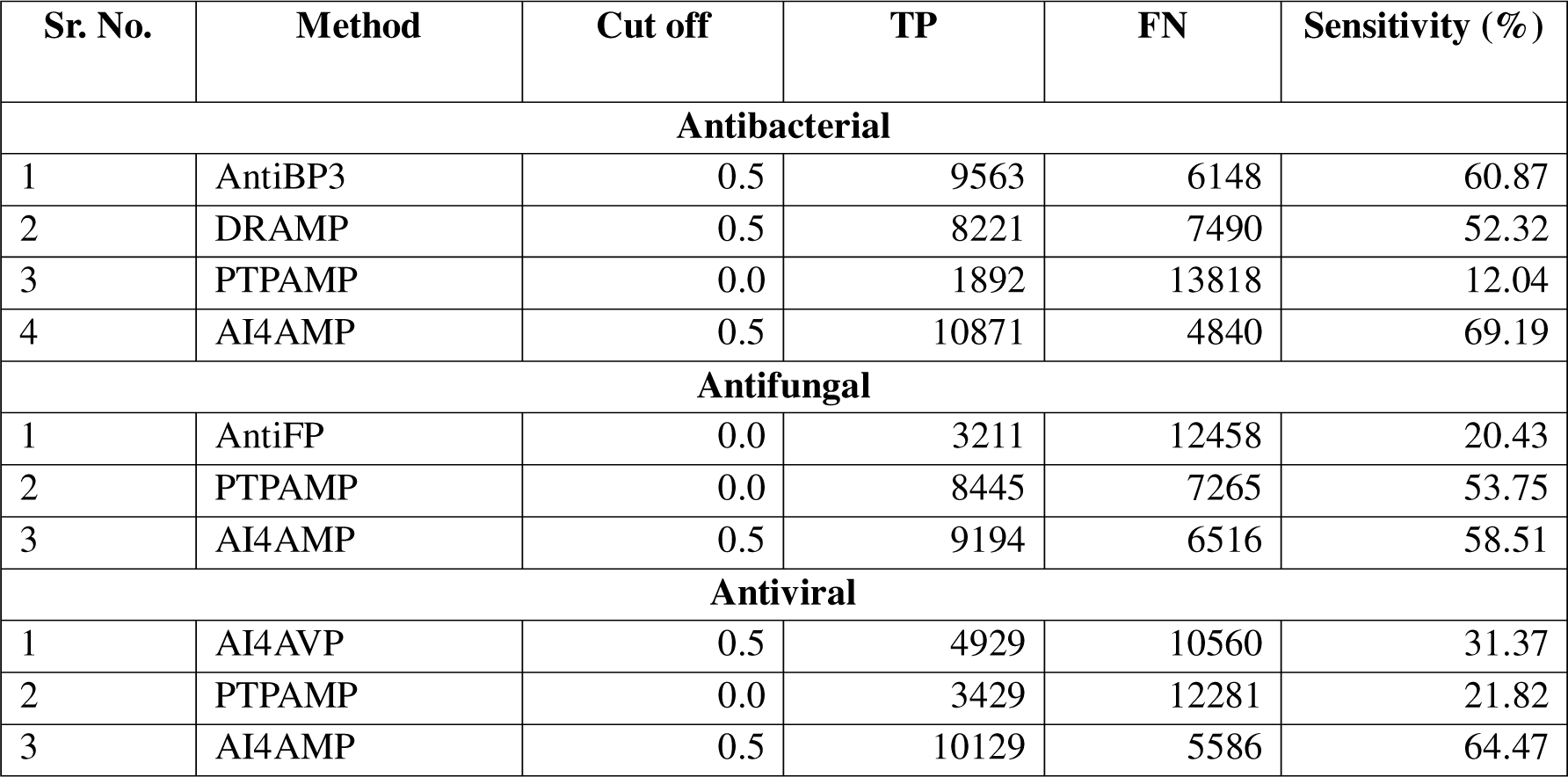
Prediction Performance of Various Methods in Classification of Peptide in a Class. Peptide sequences were provided as an input to the webservers and based on the results true positive (TP) and false negative (FN) were computed. Further they were used to compute the sensitivity (ability to predict positive class).

### Creation of refined set of peptides in each class for various analyses

To create a dataset of peptides with high confidence values, we selected peptides with high prediction scores. We set a cutoff of 0.8 or above for tools with a score range between 0 and 1. For software with different score ranges, we implemented specific cutoffs. For example, PTPAMP’s cutoff varies by peptide class: 0.5 for ABPs, 0.8 for AVPs, and 2 for AFPs. In the end, we obtained 9,175 ABPs, 6,133 AFPs, and 6,025 AVPs. The prediction scores using various in silico tools for ABPs, AVPs, and AFPs are provided in Supplementary Table S1-S3. As some peptides may be common among two or more classes, we filtered the common and class-specific peptides. As shown in Figure 1A, we identified 3,363 common peptides, 2,449 unique ABPs, 272 unique AFPs, and 1,443 unique AVPs (Supplementary Table S4). Similarly, we created a negative dataset, which included 2,052 common peptides, 2,331 unique non-ABPs, 191 unique non-AFPs, and 3,831 unique non-AVPs (see Methodology) (Supplementary Table S5) (Figure 1B).

**Fig 1.**
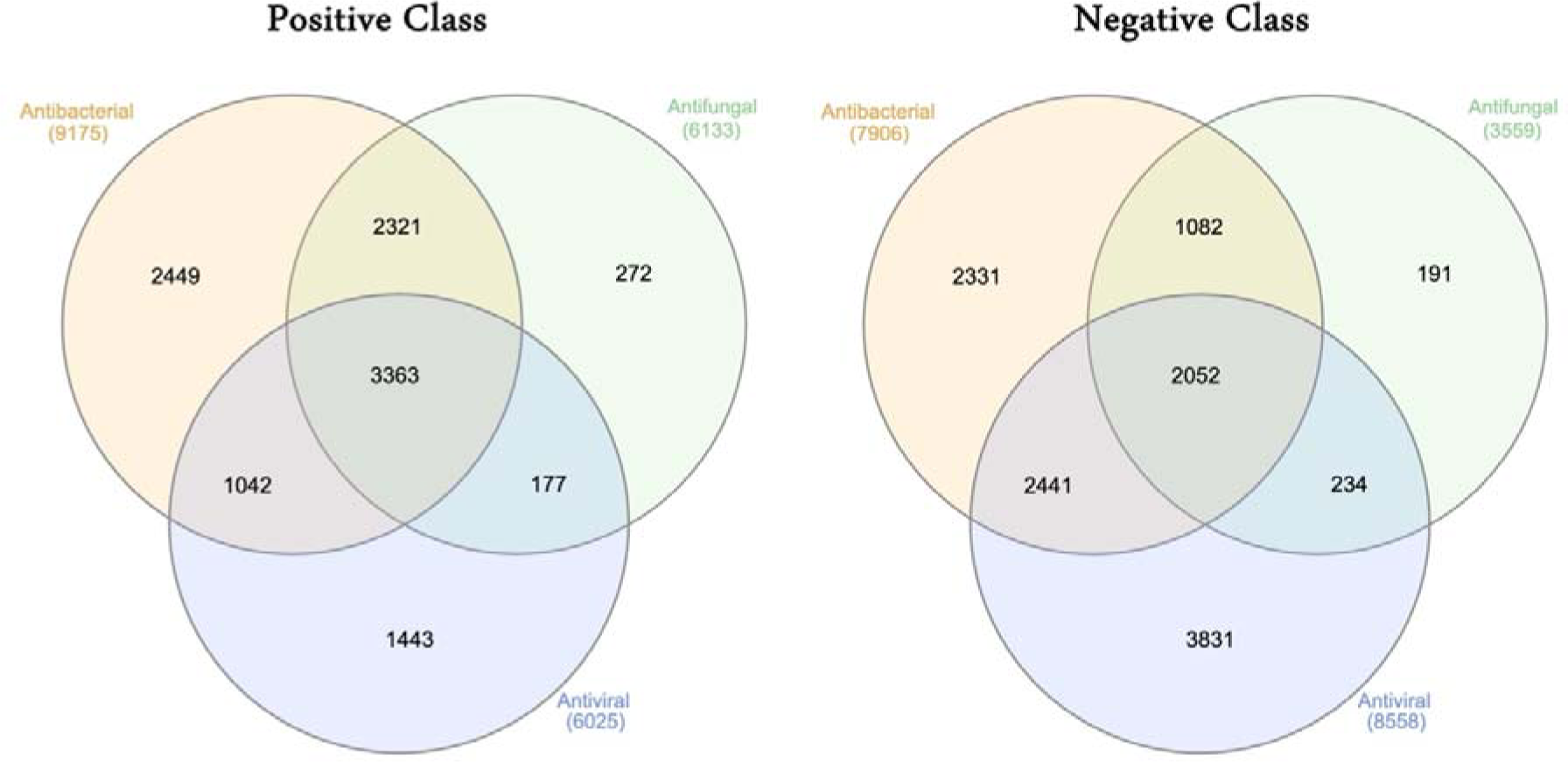
Common and Class Specific AMPs Characterization. Venn Diagram showing common and unique class specific predicted peptides in positive class (A) and negative class (B).

This dataset was designated as “High Quality Dataset 1 (HQ_DS1).” We also created “High Quality Dataset 2 (HQ_DS2)” and “High Quality Dataset 3 (HQ_DS3).” HQ_DS2 contains the same positive peptides, but for the negative dataset, we compiled data from previously published papers for each respective class. Lastly, in HQ_DS3, the positive peptides remain the same, but for negative peptides, we considered the other two positive classes as the negative set. For example, in the case of ABPs, AFPs and AVPs were considered as the negative dataset. Results for HQ_DS2 are discussed in the main paper, whereas results for HQ_DS1 and HQ_DS3 are provided as part of the supplementary.

### Amino acid composition analysis reveals enrichment of key residues in each class

We computed the amino acid composition for the three classes of AMPs. Figure 2A depicts a comparison of the average amino acid composition among ABPs, AVPs, and AFPs. We observed a high composition of Ala (A), Cys (C), Lys (K), Pro (P), and Arg (R) in ABPs. Amino acids enriched in AFPs include Asp (D), Glu (E), Phe (F), Gly (G), His (H), Asn (N), Ser (S), Thr (T), and Trp (W). Lastly, AVPs showed high enrichment of Ile (I), Leu (L), Met (M), and Val (V). We also compared the amino acid composition of each class of AMPs individually with their negative counterparts. We selected HQ_DS2 for this comparison because majority of the negative peptides in this dataset are either experimentally validated or randomly generated and have been used previously. When examining the amino acid composition of these peptides, we identified several residues that are highly enriched in the positive class compared to the negative class. In ABPs, we observed high enrichment of Lys (K), Val (V), Ala (A), Ile (I), Arg (R), Leu (L), and Met (M) (Figure 2B). In AFPs, residues such as Lys (K), Gly (G), Ile (I), Phe (F), Arg (R), Met (M), and Thr (T) were highly enriched compared to non-AFPs (Figure 2C). In AVPs, residues such as Val (V), Ile (I), Leu (L), and Met (M) were found to be enriched compared to non-AVPs (Figure 2D).

**Fig 2.**
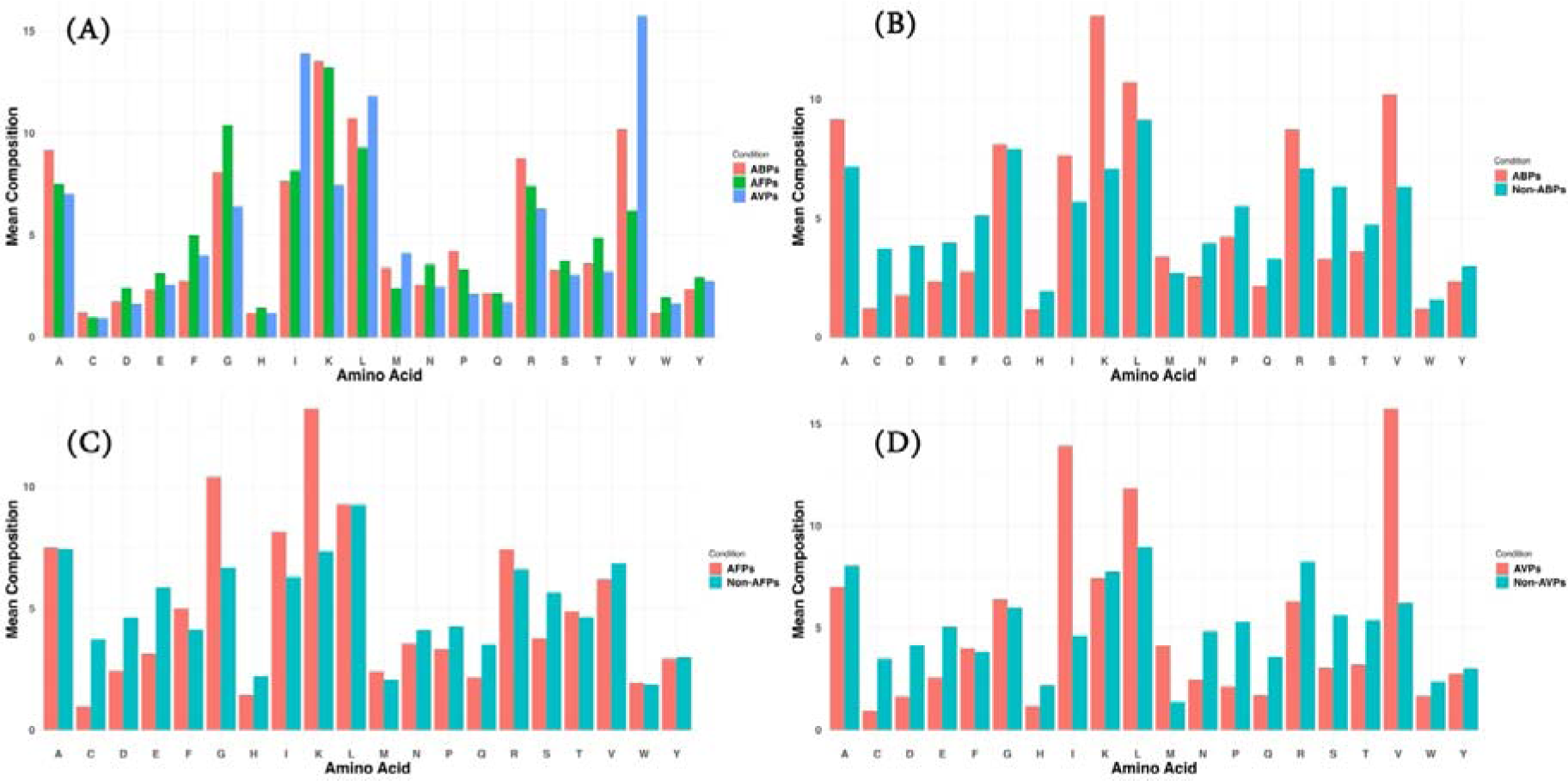
Amino Acid Composition Analysis. Average amino acid composition percentage was computed among only positive class (A); Antibacterial peptide and non-antibacterial peptides (B); Antifungal peptide and non-antifungal peptides (C); and Antiviral peptide and non-antiviral peptides present in “HQ_DS2”.

This analysis highlights that different classes of AMPs show enrichment of distinct amino acid compositions. Average amino acid composition values for the positive and negative datasets (all three) are provided in Supplementary Table S6.

### Classes of AMPs shows distinct physiochemical properties

Santos Junior et al. reported that peptides in AMPSphere are positively charged (4.7 ± 2.6) with a high isoelectric point (10.9 ± 1.2). However, we observed variation in the values when we analysed the physicochemical properties i.e., molecular weight, charge, isoelectric point, and instability index, in a class-specific manner for the peptides present in our dataset. We created boxplots of these properties (Figure 3A-D) and performed Wilcoxon tests to check if the values among the three classes are statistically significant.

**Fig 3.**
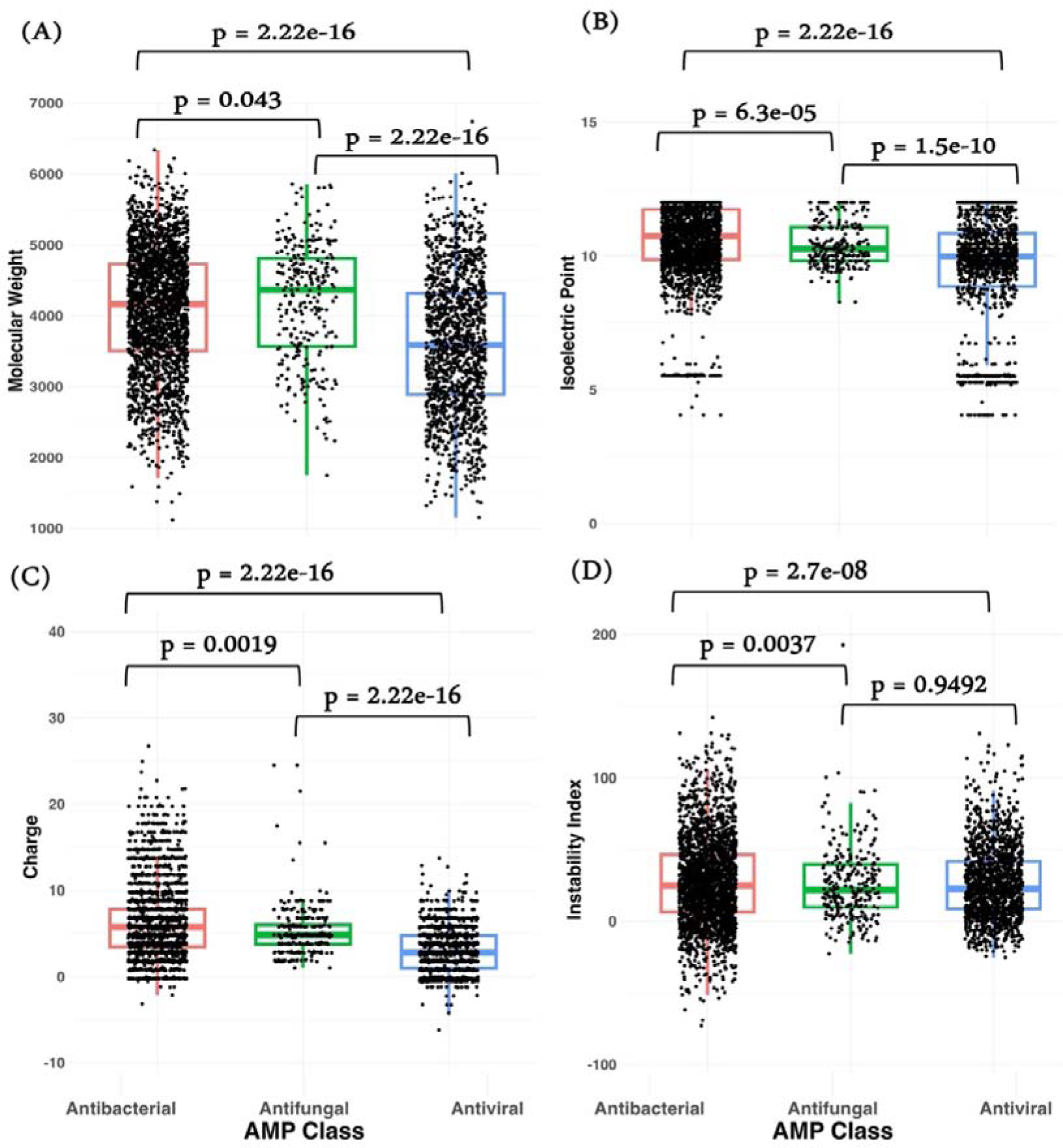
Physiochemical Properties comparison. Boxplot analysis showing distribution of Molecular Weights (A); Isoelectric Point (B); Charge (C); and Instability Index (D) of the peptides present in each class of “HQ_DS2” dataset. Mann-Whitney Test was performed to compute the statistical significance in the form of p-value.

First, we analyzed molecular weight and observed that ABPs and AVPs broadly range between 1000 and 6500, while AFPs range between 2000 and 6000. The median molecular weight of AFPs was the highest among the three classes, followed by ABPs, with AVPs having the lowest median. The differences were highly statistically significant (Figure 3A). Next, we compared the isoelectric point, finding that ABPs and AVPs range between 4 and 12, whereas AFPs range between 8 and 12. The Wilcoxon test indicated that these differences are statistically significant (Figure 3B). In terms of charge, ABPs showed the greatest variability, ranging from -3 to 27, while AVPs ranged from -10 to 13. AFPs were observed to be only positively charged, ranging from 1 to 25. These values were also found to be statistically significant (Figure 3C). Lastly, we analyzed the instability index of these peptides. ABPs exhibited the highest variation, with values ranging from -72 to 292, followed by AFPs ranging from -22 to 193, and AVPs ranging from -25 to 130. The Wilcoxon test revealed that these differences are statistically significant (Figure 3D). Average value of physicochemical properties of ABPs, AFPs and AVPs are provided in Supplementary Table S7.

### Residue Positional Preference analysis

We conducted positioning analyses for all three classes using all three sets. Here, we explain the results using the peptides from the positive class and their respective negative datasets. We created Two Sample Logos (TSLs) using the first 10 residues from the N-terminus of each peptide.

For ABPs, we observed a preference for the positively charged amino acid Lys (K), followed by Ala (A) and Arg (R) at the first position. At other positions, Val (V) and Lys (K) were dominant along with Gly (G), Ile (I), and Met (M) (Figure 4A). In AFPs, we observed a high enrichment of Leu (L), followed by Ile (I) and Val (V) at the first position. At the second position, Ala (A) was dominant, followed by Ile (I) and Gly (G). At the third position, Gly (G) was the most preferred residue, followed by Phe (F) (Figure 4B). Lastly, for AVPs, we observed enrichment of Leu (L), Ile (I), Val (V), and Met (M) at the first position. At the second and third positions, there was enrichment of Ile (I) and Val (V) along with Gly (G) and Ala (A) (Figure 4C).

**Fig 4.**
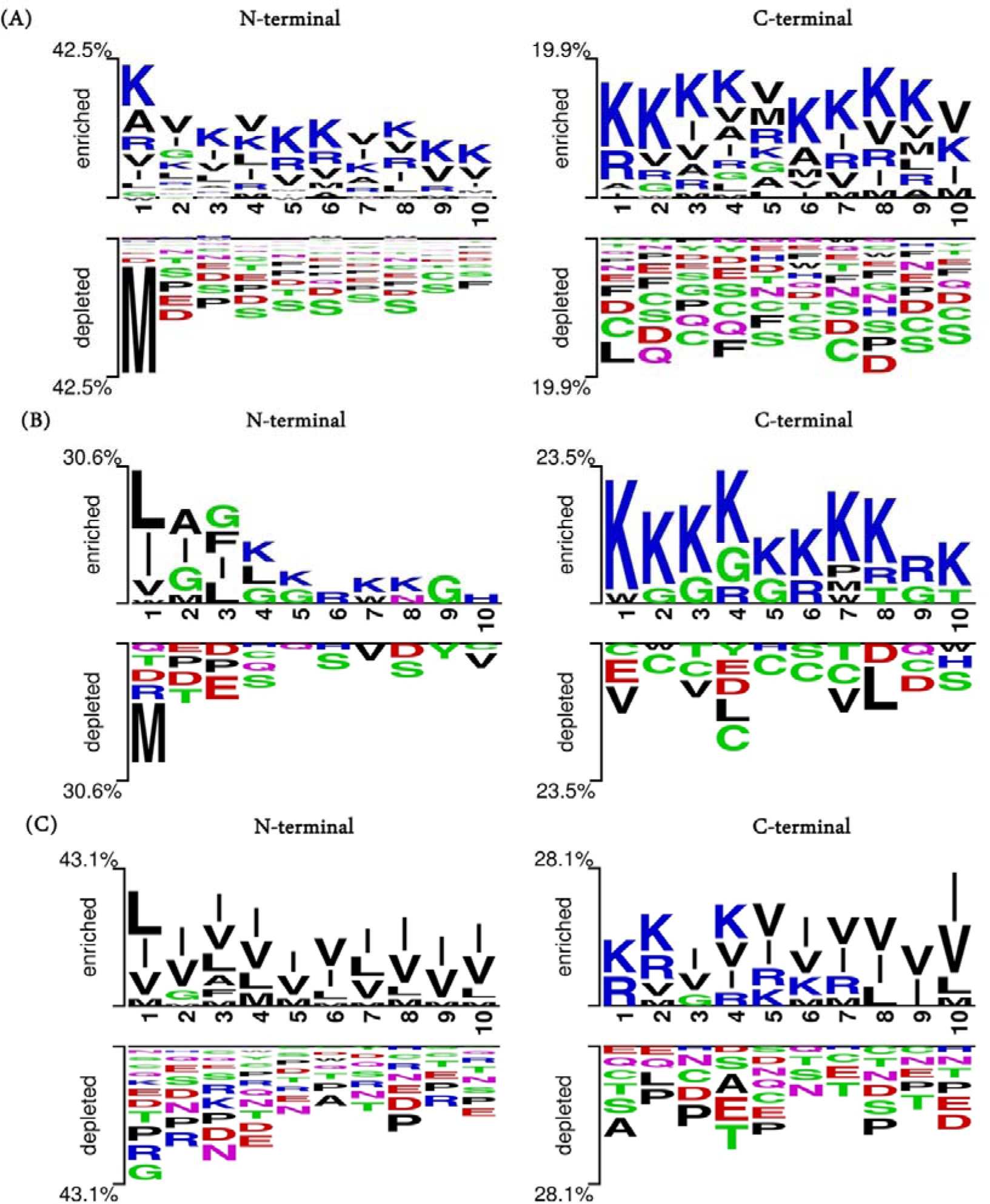
Positional Preference Analysis. Two Sample Logo analysis showing the amino acid residue preference for the first 10 residues from N-terminal and C-terminal in ABPs and non-ABPs (A); AFPs and non-AFPs (B); and AVPs and non-AVPs (C); present in “HQ_DS2”.

We also analyzed the positional preference of residues at the C-terminus, creating TSLs using the first 10 residues for all three datasets. In the dataset with Pos and “Neg_DS2,” we observed the dominance of Lys (K) for the first three positions, followed by Arg (R) and Val (V). Gly (G) was also preferred at various positions, such as the second, fourth, and fifth (Figure 4A). Similarly, in AFPs, Lys (K) and Trp (W) were dominant at the first position. At the remaining positions, Gly (G) was the second preferred residue after Lys (K) (Figure 4B). Lastly, in AVPs, Lys (K) and Arg (R) were preferred at the first and second positions, while Ile (I) and Val (V) were preferred at other positions (Figure 4C).

The current analysis shows the diversity of residues at the N- and C-termini in each class, highlighting the different biology and mechanisms of action of these peptides. Positional preferences for all three classes of peptides associated with datasets HQ_DS1 and HQ_DS3 are shown in Supplementary Figures S1 and S2, respectively.

### Motif Analysis

Exclusive motifs were characterized from positive and negative peptides in each class using MERCI software. The results showed that ABPs were enriched for motifs such as “VRVR,” “RVRVR,” and “AKKPA,” while non-ABPs were enriched for motifs such as “HGFR,” “MKRT,” and “KHKQ.” In the case of AFPs, we observed enrichment of motifs such as “DFFAI,” “FIVFTI,” and “KFRR,” whereas non-AFPs were enriched for motifs such as “CP,” “DCP,” “LCE,” and “GCN.” AVPs showed enrichment of motifs including “VVV,” “VLV,” and “IIII,” while non-AVPs showed enrichment of motifs such as “DRLN,” “PCR,” and “RLNE.”

A complete list of exclusive motifs in positive and negative set for all the 3 classes are provided in Supplementary Table S8. Similarly, we also identified the motifs present in the other two datasets (HQ_DS1 and HQ_DS3), with complete lists provided in Supplementary Tables S9 and S10, respectively.

### Machine Learning Model Development and Performance

In this study, we developed and evaluated a range of machine learning classifiers using HQ_DS2 dataset to predict anti-bacterial, anti-fungal, and anti-viral properties of the peptides. These predictions were based on various composition-based features generated using the Pfeature tool. The dataset was divided into training (80% of the data) and validation set (20% of the data) and five-fold cross validation technique was implemented. Performance was computed in both threshold dependent and threshold independent manner (See Methods). The performance of each classifier across the different categories such as, anti-bacterial, anti-fungal, and anti-viral, is detailed in Supplementary Tables S11, S12, and S13, respectively.

Our analysis showed that models using amino acid composition (AAC) with the Extratrees (ET) classifier performed comparably to the top-performing models based on more complex compositional features. Although AAC was not the single best-performing feature set, its results were on par with those of other leading features. Notably, the AAC feature set is the smallest, consisting of only 20 features, which presents a significant advantage in machine learning. This compact size allows for the development of models that are not only efficient but also easier to interpret and implement, without sacrificing predictive accuracy. This balance of simplicity and performance underscores the practical value of AAC as a feature set for predicting the antimicrobial potential of peptides.

As shown in Table 2, for the classification of anti-bacterial versus non-anti-bacterial peptides, the extra trees-based model achieved the highest AUROC of 0.983 on the training dataset and 0.984 on the independent dataset. In the case of anti-fungal versus non-anti-fungal peptides, the AAC-based extra trees model reached the highest AUROC of 0.987 on the training dataset and 0.986 on the independent dataset. Similarly, for antiviral versus non-antiviral peptides, the highest performance was achieved with the extra trees model, recording an AUROC of 0.988 on the training dataset and 0.989 on the independent dataset.

**Table 2:**
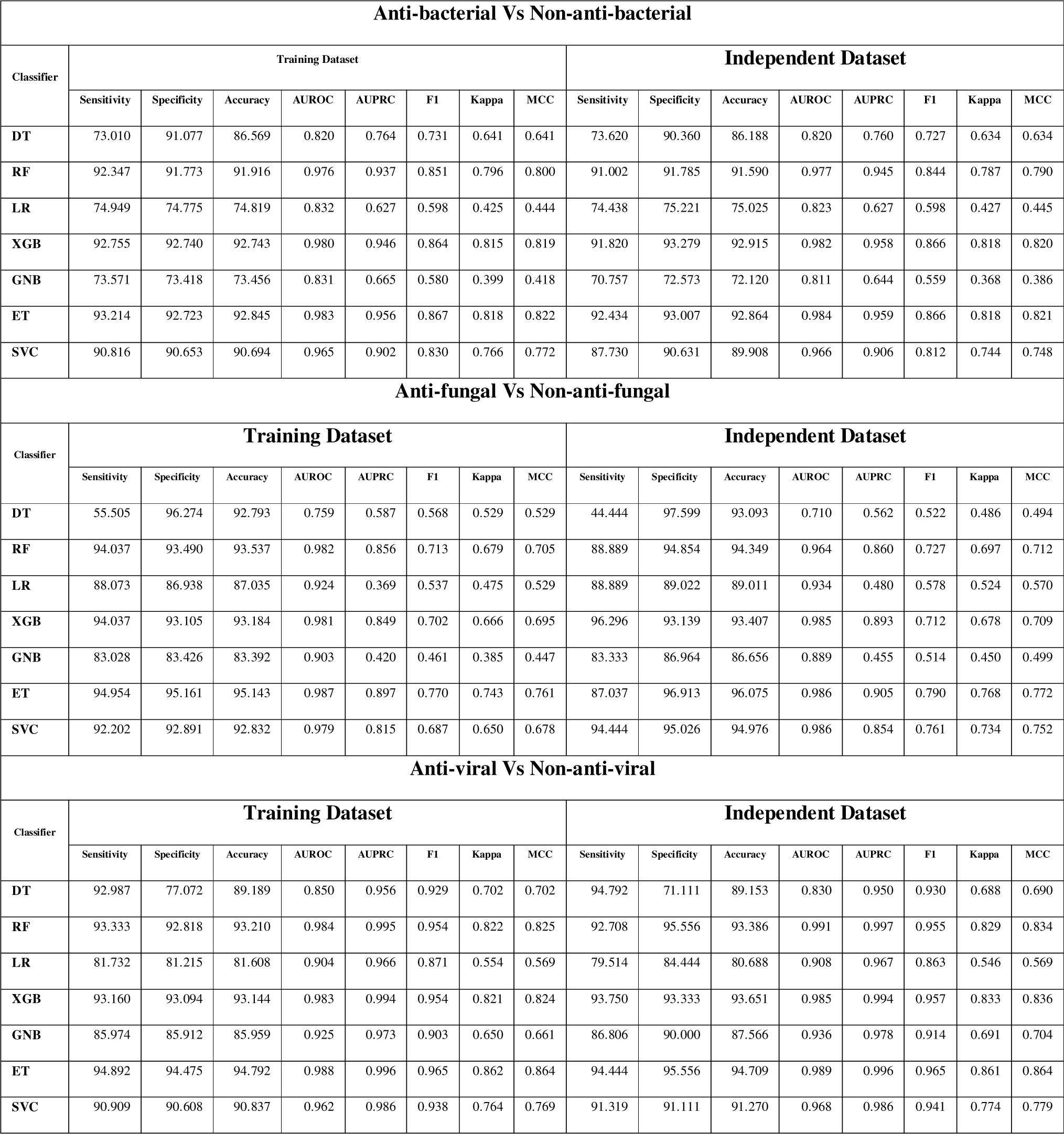
Machine learning models classify peptides with high accuracy. Performance of seven distinct classifiers on training and independent dataset for predicting anti-bacterial, anti-fungal, and anti-viral peptides. Both threshold dependent and independent measures were computed to evaluate the model performance.

### Identification of peptides with therapeutic potential

An ideal peptide candidate with therapeutic potential should possess cell-penetrating properties and should not be toxic and allergic. We screened the ABPs, AVPs, and AFPs using various recently developed in silico tools and identified the peptides fulfilling all the three criteria. We used the ToxinPred 3.0 server to predict toxicity, AlgPred 2.0 for predicting allergenicity and CellPPD for predicting cell-penetrating potential. In the end, we identified 322 ABPs, 29 AFPs, and 102 AVPs (Supplementary Table S14). Next, we predicted the secondary structure of these peptides using the GOR4 webserver, as secondary structure provides important insights into the peptide mechanism of action. For example, α-helix is commonly found in AMPs and is crucial for their ability to insert into and disrupt microbial membranes [43]. Likewise, amphipathic β-sheets are another common pattern in AMPs, facilitating the insertion of peptides into the microbial membrane lipid bilayer [44]. Additionally, a small portion of AMPs adopts random coil structures, exhibiting flexibility to interact with multiple targets [43]. Full output using GOR4 for each class is provided in Supplementary Table S15-S17.

As shown in the violin plot, the alpha-helix was predominantly present in ABPs, with a median content of 44.09%, followed by AVPs and AFPs, with median values of 27.91% and 21.52%, respectively. The Wilcoxon test showed significance between ABPs & AFPs and AFPs & AVPs but not between AFPs and AVPs (Figure 5A). Similarly, beta sheets were predominantly found in AFPs, with a median value of 41%, followed by AVPs with 32%, and ABPs with 16%. The Wilcoxon test showed significant differences between ABPs & AFPs and AFPs & AVPs, but not between ABPs and AVPs (Figure 5B). However, random coils were found to be equally distributed among the three classes. The median percentage of random coils was 35% in ABPs and AVPs, and 25% in AFPs. The Wilcoxon test showed a significant difference between ABPs and AVPs (Figure 5C).

**Figure 5.**
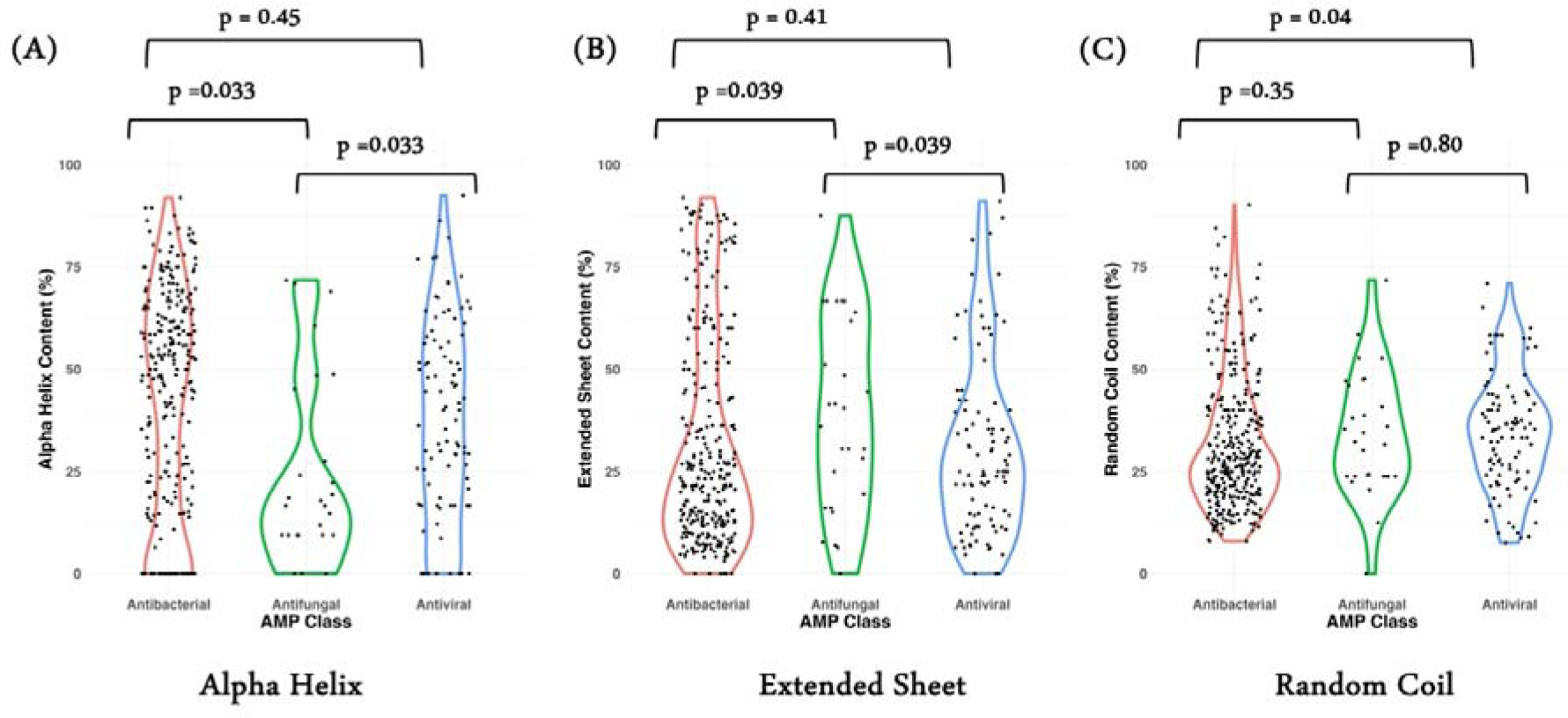
Secondary Structure Distribution in 3 class of AMPs. Violin Plot showing percentage distribution of Alpha helix (A); Extended Sheet (B); and random Coil (C) computed using GOR4 server in each class of peptides. Mann-Whitney Test was performed to compute the statistical significance in the form of p-value.

Lastly, we selected the top 10 peptides with the highest half-life in blood for each class, i.e., ABPs, AFPs, and AVPs, which may possess high therapeutic properties (Table 3). Half-life results all the peptides in each class is provided in Supplementary Table S18.

**Table 3.**
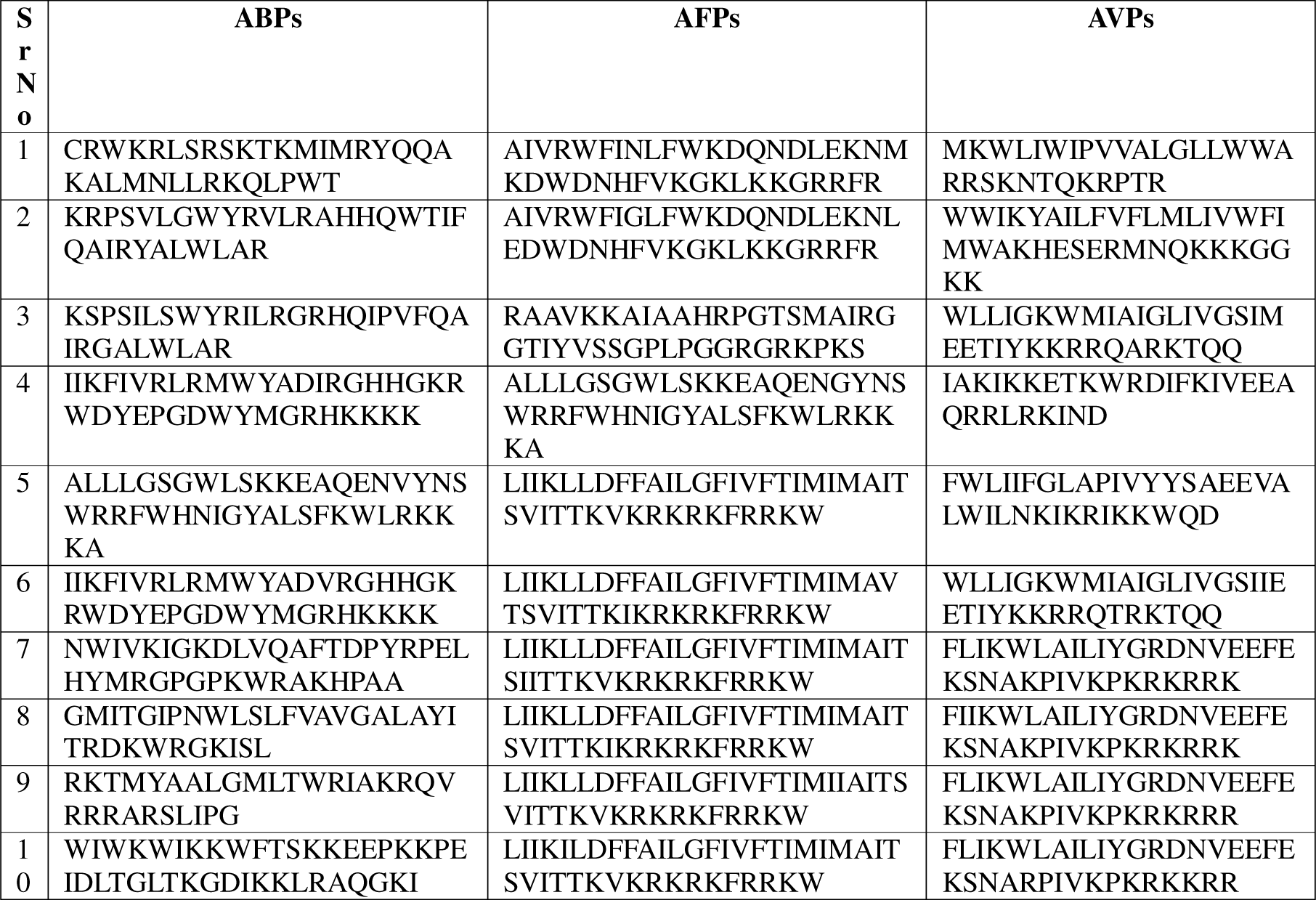
List of Top10 Peptides with Therapeutic Potential. Amino acid Sequence of top10 peptides selected in each category based on multiple in silico studies followed by maximum half-lives.

### Peptide Structure Prediction and Molecular Docking

ColabFold embedded in ChimeraX1.8 software was used for predicting the structure of the top10 peptides (Table 2) selected in each class of AMPs. Confidence of the predicted structures is measured in two parameters (i) pLDDT score and (ii) MSA. pLDDT plots, show the predicted local distance difference test scores for each residue position Regions with a pLDDT over 90 are expected to be accurately modelled, which suggests they can be used for accurate model-based applications. Other regions with a pLDDT > 70 have similar levels of model uncertainty. Conversely, whether you should take regions with a pLDDT score between 50 and 70 seriously (low confidence) varies reciprocally. The 3D structures of regions with pLDDT scores below 50 are often seen to take on a single, ribbon-like character and should not be considered reliable. Supplementary Figure S3 (a-c) represent the pLDDT plots for each of the 5 generated models of the top10 ABPs, AFPs and AVPs. The MSA is represented as a heat map in which each sequence has its own alignment to the input matrix, leading to corresponding columns of the rows–columns cell. The color gradient indicates the level of identity scored in response, with sequences from top to bottom ranked according to their levels, i.e., highest peak at the head and lower below (Supplementary Figure S4(a-c)). White regions showed areas not covered, which is a result of subsequence entries in the database. The black line indicates the fraction of sequences in which that position is aligned. The pLDDT and pTM scores alongside the peptide sequences are provided in Supplementary Tables S19-S21 for ABPs, AFPs and AVPs, respectively.

Next, we performed molecular docking studies using the top 10 predicted peptides with case example in each class of AMPs. As a case study, we selected one pathogenic protein from an organism representing a class of AMPs. We used HDOCK webserver and default parameters to perform the docking analysis. The best docked posed complexes of ABPs with Streptococcus pneumoniae protein (PDB Id: 3ZPP), for AFPs with Candida albicans protein (PDB Id: 4LE8), and for AVPs with SARS-COV2 protein (PDB Id: 6LU7) are represented in Figures 6(a), 7(a), and 8(a), respectively and representation of all the interacting residues for each of the best docked posed complexes is provided in Figures 6(b), 7(b), and 8(b), respectively.

**Figure 6(a).**
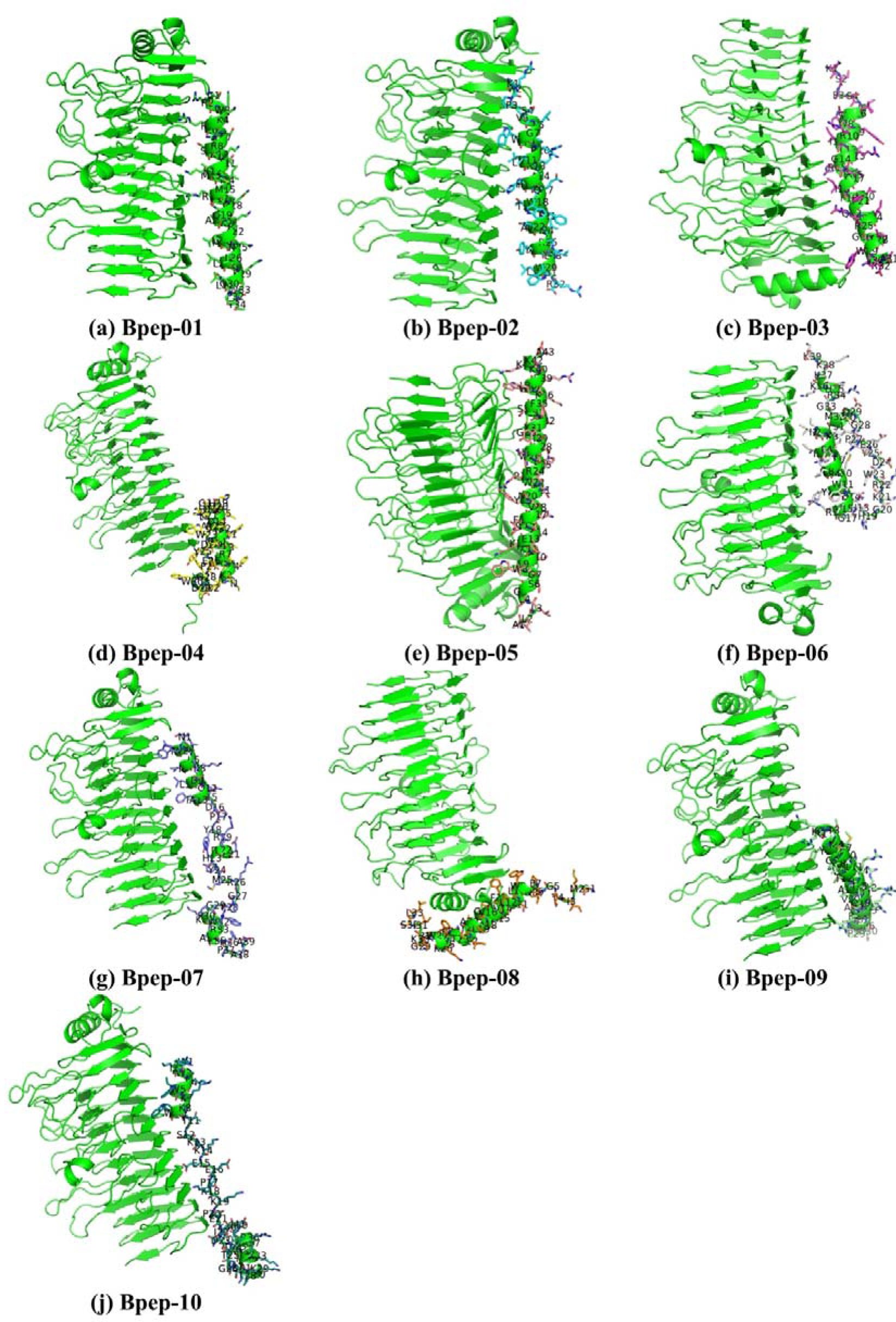
The 3D docked complexes (a-j) of the alpha fold predicted anti-bacterial peptides with Streptococcus pneumoniae protein (PDB Id: 3ZPP). Figure (a-j) represents one letter code for each of the peptide residues

**Figure 6(b).**
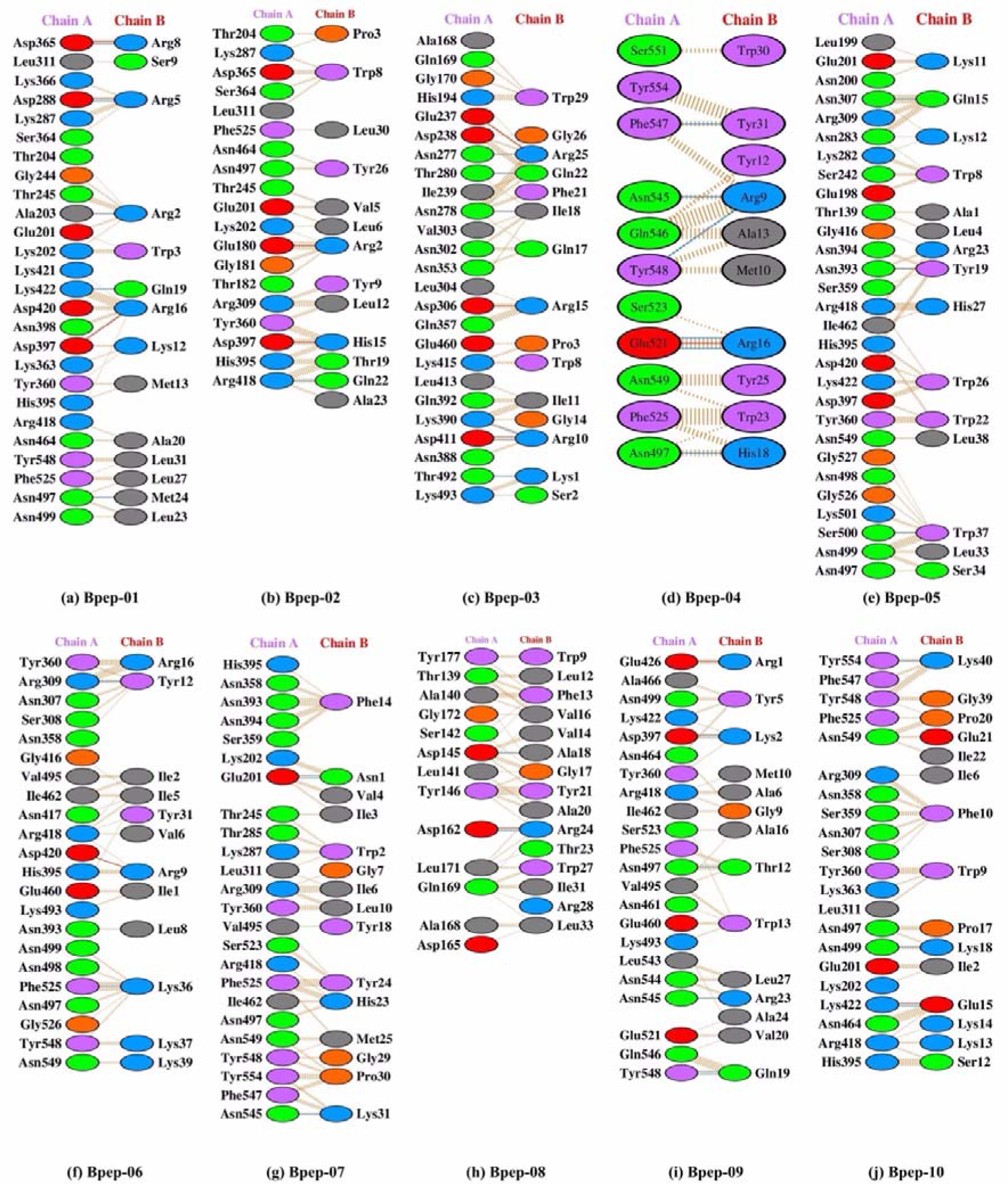
The interacting residues in the docked complexes of the alpha fold predicted anti-bacterial peptides with Streptococcus pneumoniae protein (PDB Id: 3ZPP). Chain-A represents the Streptococcus Pneumoniae protein residues whereas chain-B represents the alpha fold predicted anti-bacterial peptide residues.

**Figure 7(a).**
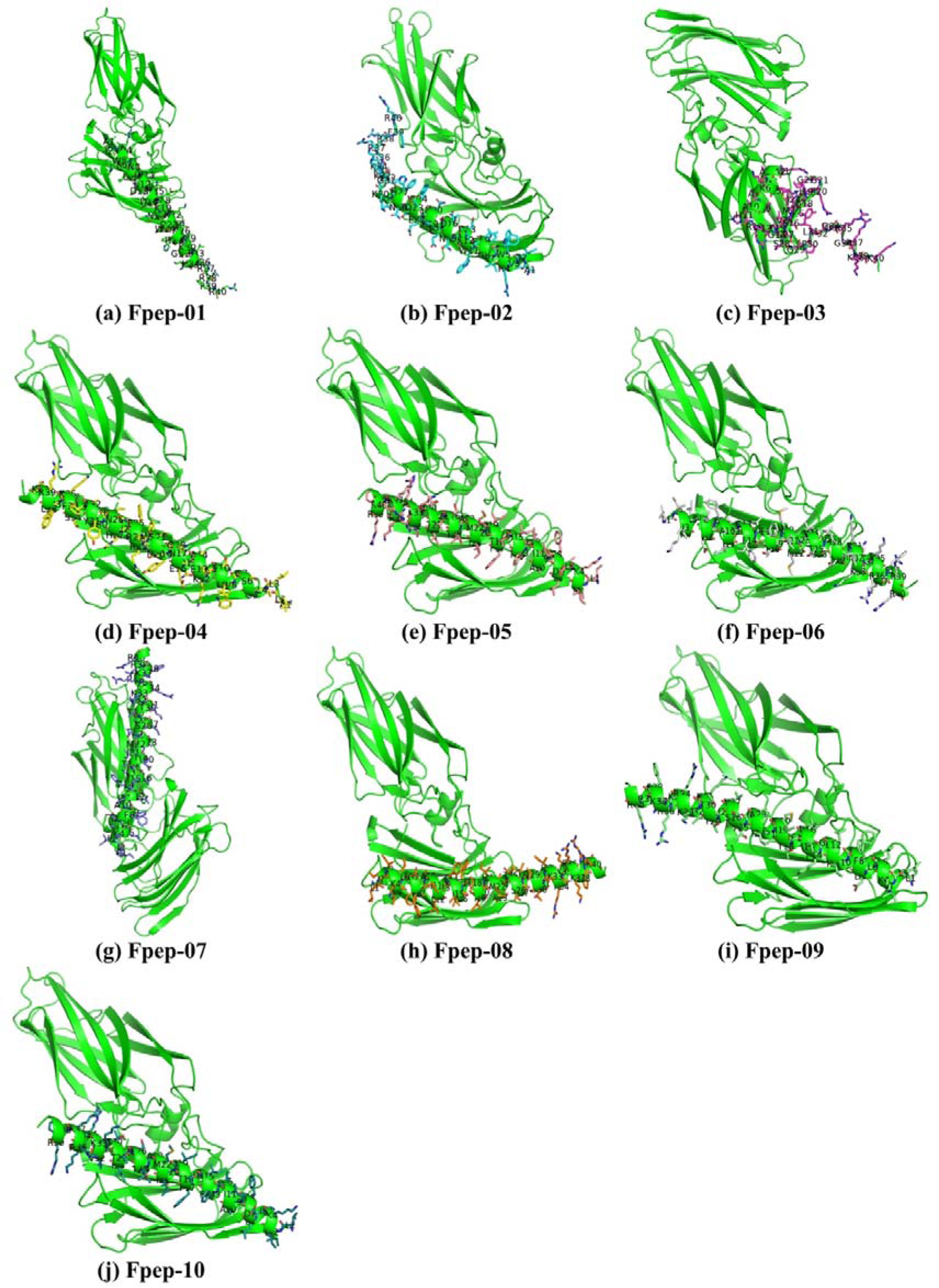
The 3D docked complexes (a-j) of the alpha fold predicted anti-fungal peptides with Candida albicans protein (PDB Id: 4LE8). Figure (a-j) represents one letter code for each of the peptide residues.

**Figure 7(b).**
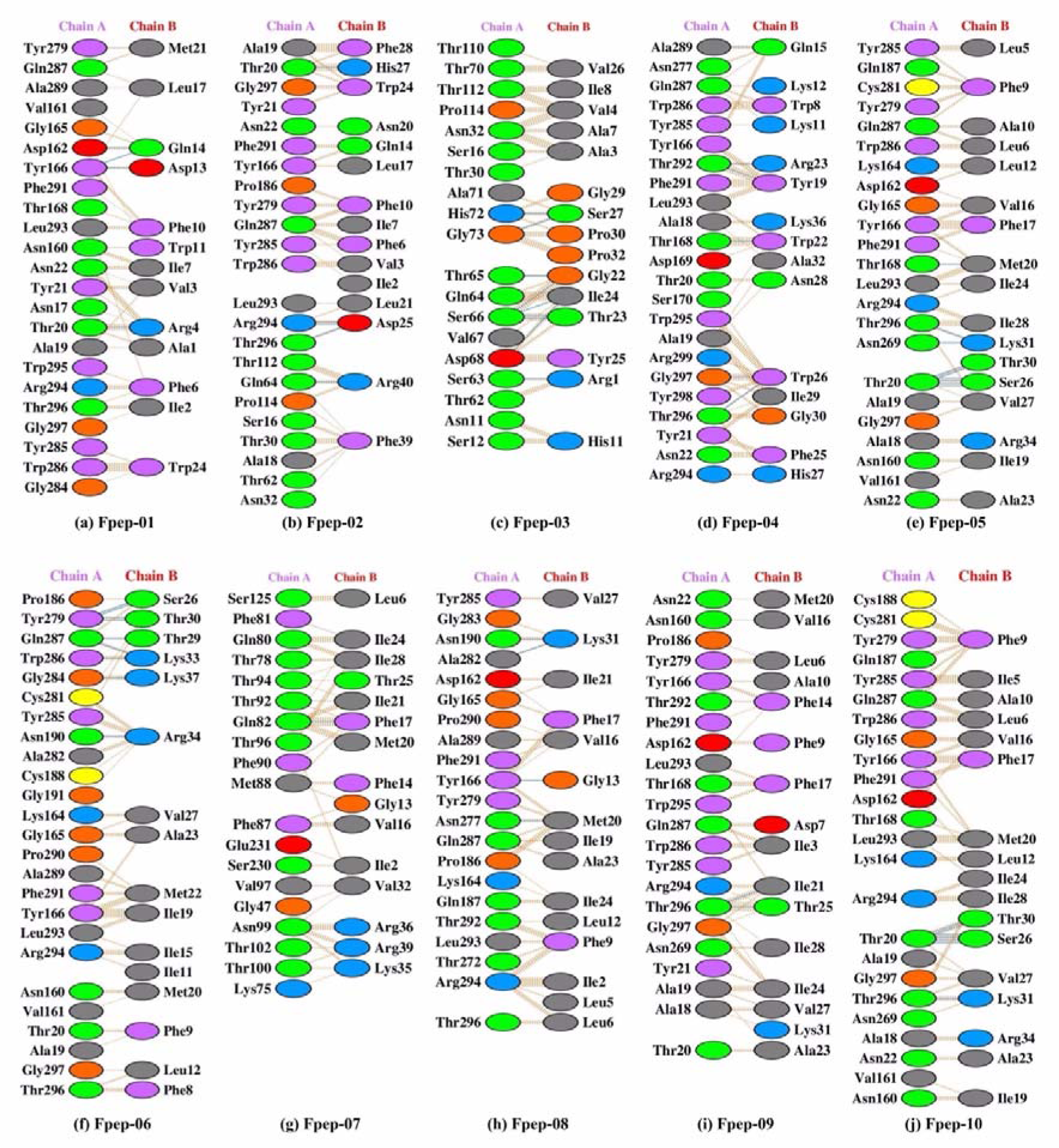
The interacting residues in the docked complexes of the alpha fold predicted anti-fungal peptides with Candida albicans protein (PDB Id: 4LE8). Chain-A represents the Candida Albicans protein residues whereas chain-B represents the alpha fold predicted anti-fungal peptide residues.

**Figure 8(a).**
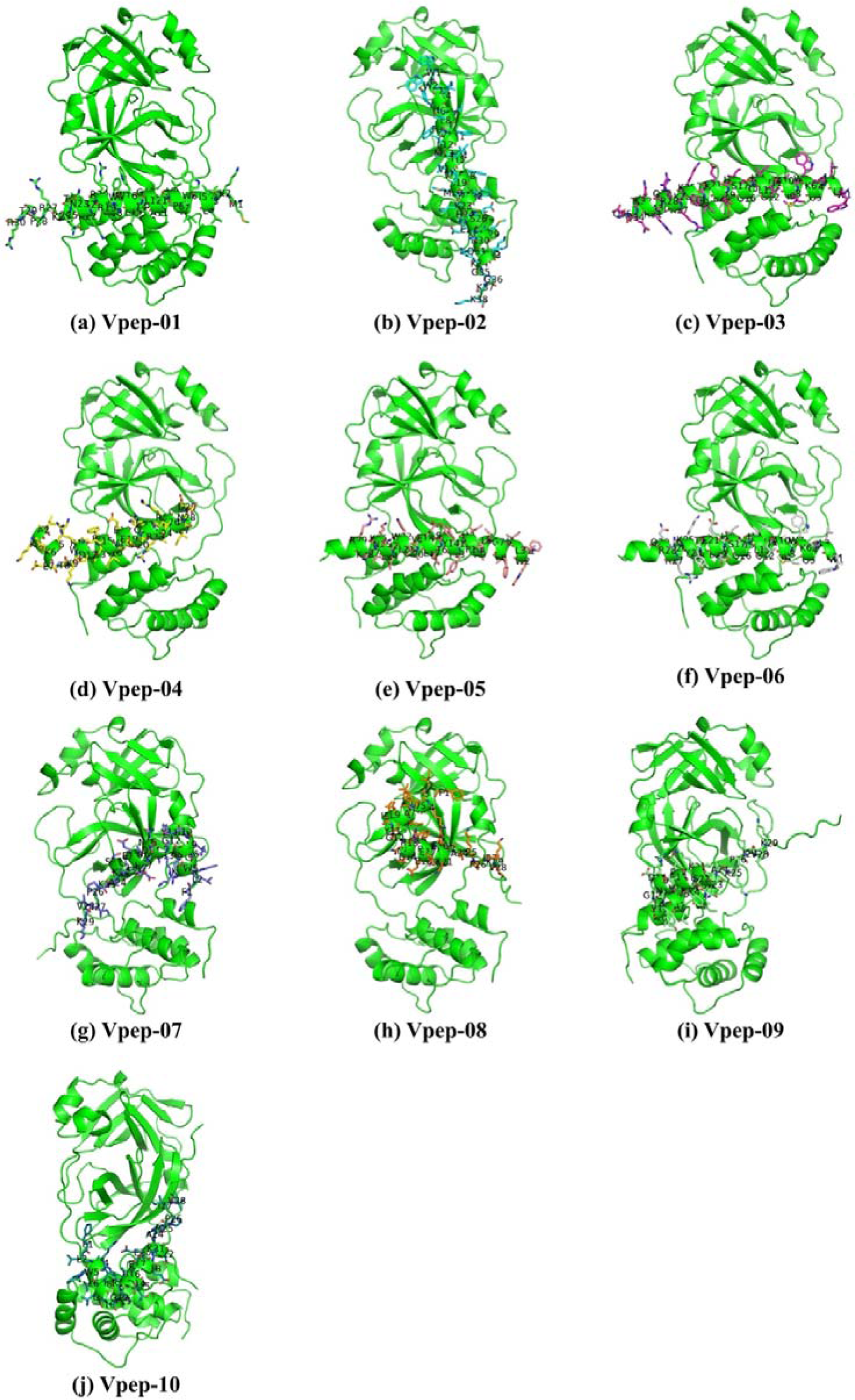
The 3D docked complexes (a-j) of the alpha fold predicted anti-viral peptides with SARS-COV-2 viral protein (PDB Id: 6LU7). Figure (a-j) represents one letter code for each of the peptide residues.

**Figure 8(b).**
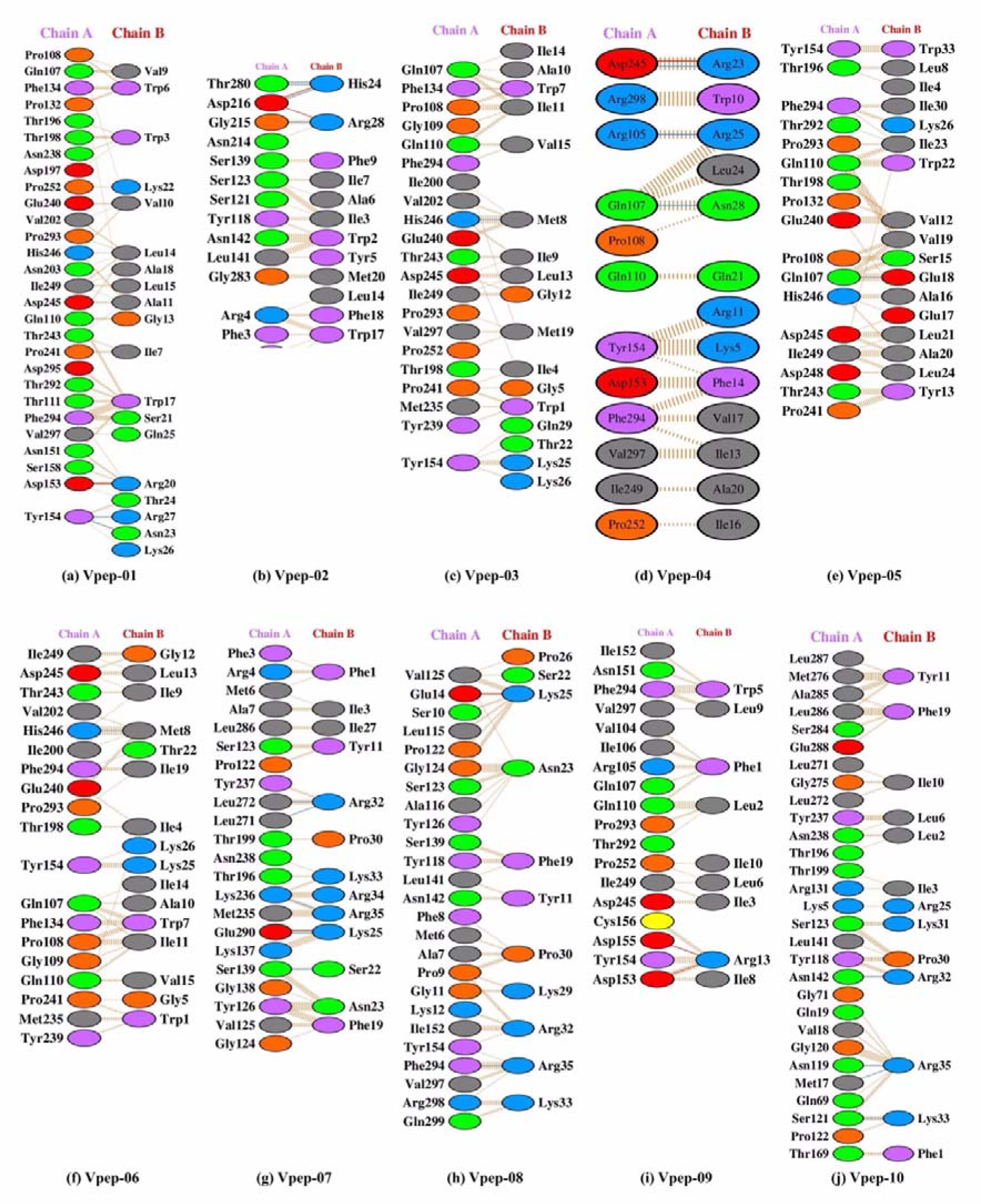
The interacting residues in the docked complexes of the alpha fold predicted anti-viral peptides with SARS-COV-2 viral protein (PDB Id: 6LU7). Chain-A represents the SARS-COV2 viral protein residues whereas chain-B represents the alpha fold predicted viral peptide residue.

A cursory view of figure 6(a), 7(a), and 8(a), gives us an idea that in general the peptides tend to orient itself in such a direction so as to complement the protein by exposing maximum of its surface area, allowing a greater extent for interactions. However, in a few cases, the observed orientations are not straightforward, viz., Bpep-04, Bpep-08; Fpep-01, Fpep-03; and Vpep-07, Vpep-08, Vpep-09, and Vpep-10; wherein the peptide is either oriented in some unusual orientation or folded itself so as expose itself for maximum interactions with the peptide. When we look deep into the interacting residues, we observe that, the interactions, in general are predominantly non-covalent in nature. Observed interactions can be classified into either electrostatic (involving charged residues, such as, Asp, Glu, Lys, and Arg), cation-π (involving cationic residues, such as, Lys, and Arg along-with Phe, Trp, and Tyr), or anion-π (involving anionic residues, such as, Asp, and Glu along-with Phe, Trp, and Tyr) type of interactions, respectively. However, at some instances, some π-π type of weak interactions were also observed (involving interactions among the residues having π_e_, *viz.,* Phe, Trp, and Tyr). ABPs, in general, offer their cationic residues (Lys, Arg) or residues having π_e_ rings (Phe,, Trp, Tyr) for interactions towards the protein. AFPs on contrary offer, all type of residues, viz., cationic residues (Lys, Arg), anionic (Asp) or residues having π_e_ rings (Phe, Trp, Tyr) for interactions towards the protein. Similarly, the AVPs also offer their cationic residues (Lys, Arg) or residues having π_e_ rings (Phe, Trp, Tyr) for interactions towards the protein

In particular, in the case of ABPs, Bpep-1, Bpep-3, Bpep-5, and Bpep-10 are driven by cation-π, and electrostatic interactions, Bpep-2, and Bpep-3 are driven by anion-π, and electrostatic interactions, whereas Bpep-5, Bpep-6, Bpep-7, and Bpep-8 are predominantly driven by π-π interactions, among their residues. Further, among the AFPs, Fpep-6, Fpep-8, and Fpep-10 are driven by cation-π type of interactions, Fpep-3, Fpep-4, Fpep-5, Fpep-9, and Fpep-10 are driven by anion-π type of interactions, whereas, Fpep-1, Fpep-2, Fpep-4, Fpep-5, Fpep-8, Fpep-9, and Fpep-10 are predominantly interacting via π-π interactions. Along with these, for Fpep-3, Fpep-7, and Fpep-10, electrostatic interactions also play a part. Furthermore, in the ensemble of AVPs, Vpep-2, Vpep-3, Vpep-5, Vpep-6, Vpep-8, and Vpep-10 are governed by cation-π interactions, Vpep-1, Vpep-2, Vpep-4, and Vpep-5 are driven by anion-π interactions, all of the docked complexes of anti-viral peptides exhibit π-π interactions, and at some instances we also observed some electrostatic interactions as well, viz., Vpep-1, Vpep-3, Vpep-4, Vpep-7, Vpep-8, and Vpep-10.

The docking scores of the best docked posed complexes for the ABPs are provided in Table 4. A perusal of this table gets us to know that Bpep-05, and Bpep-07 have the highest docking scores among all the modelled ABPs, viz., -277.95 kcal/mol, and -272.53 kcal/mol, respectively, whereas Bpep-10 shows the lowest docking score of -215.28 kcal/mol. The average docking score of the ABPs is -247.79 kcal/mol; and five out of the ten modelled ABPs have their docking scores higher than the average value. Likewise, the docking scores of the best docked posed complexes of the AFPs is given in Table 5. A perusal of this tables gives us an idea that Fpep-4, and Fpep-10 exhibit the highest docking scores of -312.33 kcal/mol, and -266.74 kcal/mol, respectively, whereas, Fpep-3 has the lowest docking score of -220.64 kcal/mol. The average docking scores of the AFPs is -260.44 kcal/mol; and four out of ten modelled AFPs have their docking scores higher than the average value. Lastly, the docking scores of the best docked posed complexes of the AVPs is given in Table 6. A perusal of this tables gives us an idea that Vpep-1, and Vpep-2 exhibit the highest docking scores of -286.60 kcal/mol, and -262.46 kcal/mol, respectively, whereas, Vpep-9 has the lowest docking score of -219.82 kcal/mol. The average docking scores of the AVPs is -239.38 kcal/mol; and only top two out of the ten modelled AVPs have their docking scores higher than the average value. In general, we can state that AFPs exhibit higher docking scores, followed by ABPs, whereas the AVPs have the least docking scores. The docking scores of all the 10 docked posed complexes for the ABPs, AFPs and AVPs are provided in Supplementary Table S22-S24 respectively.

**Table 4.**
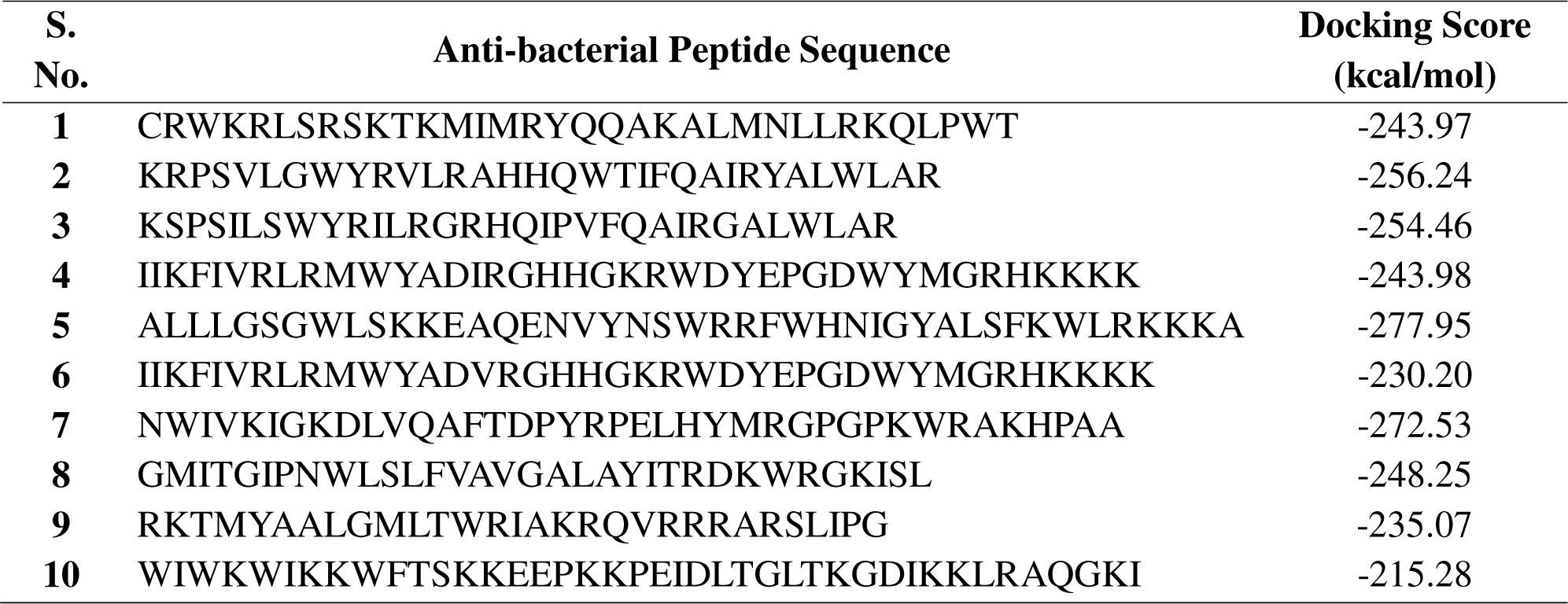
Molecular Docking analysis reveals strong binding between protein and ABPs. The peptide sequences and the docking scores of the alpha fold predicted ABPs with Streptococcus pneumoniae protein (PDB Id: 3ZPP).

**Table 5.**
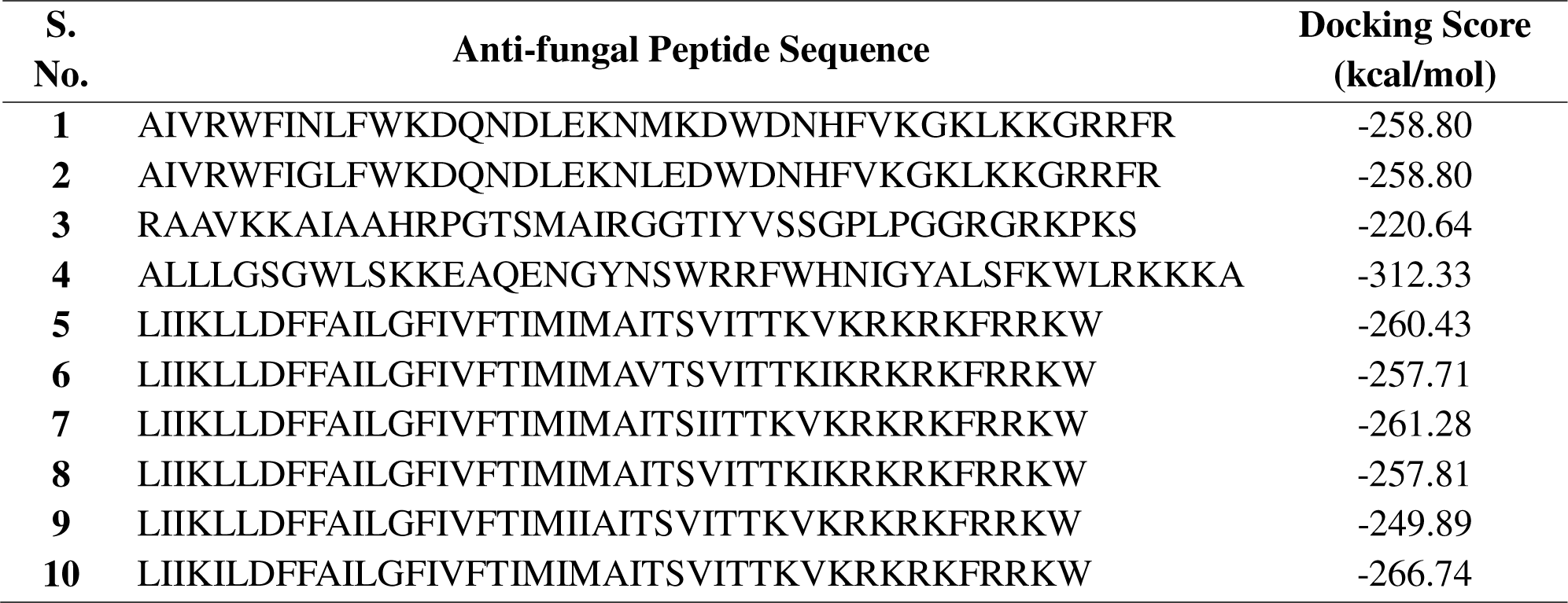
Molecular Docking analysis reveals strong binding between protein and AFPs. The peptide sequences and the docking scores of the alpha fold predicted AFPs with Candida albicans protein (PDB Id: 4LE8).

**Table 6.**
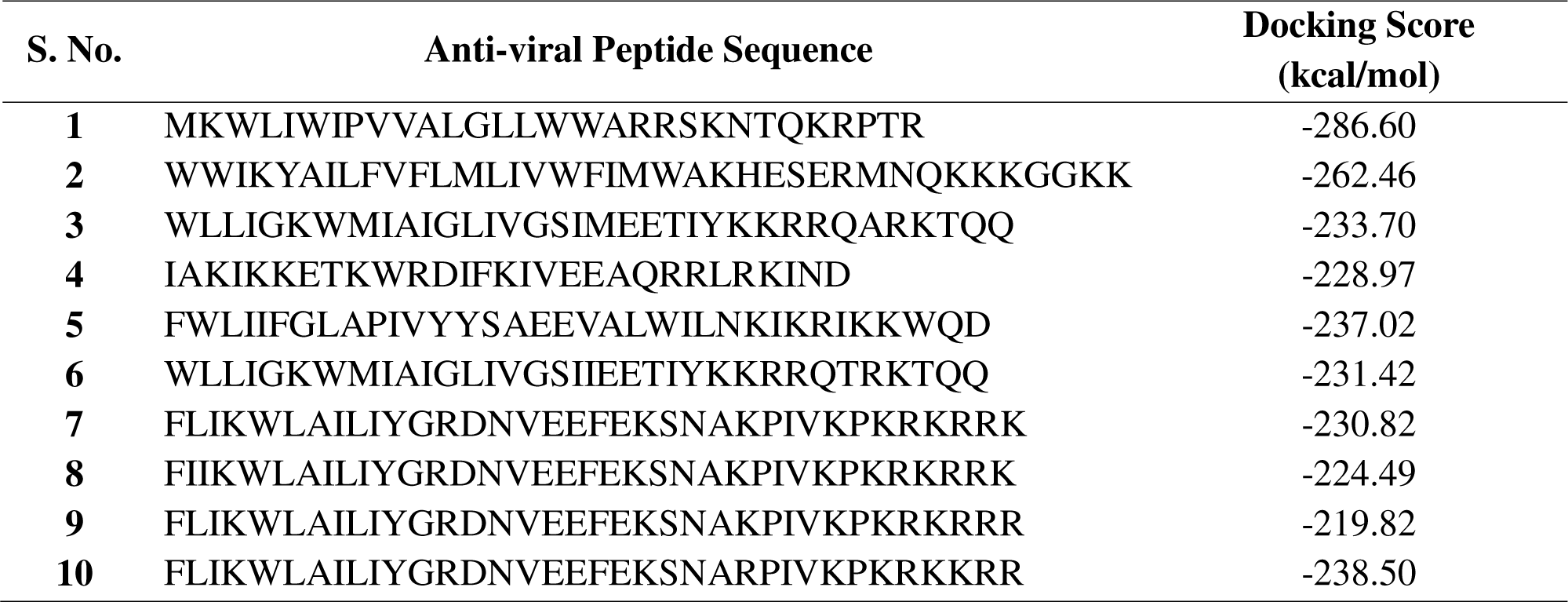
Molecular Docking analysis reveals strong binding between protein and AVPs. The peptide sequences and the docking scores of the alpha fold predicted AVPs with SARS-COV-2 viral protein (PDB Id: 6LU7).

## Discussion

Increasing antimicrobial resistance (AMR) is one of the leading global public health threats. A recent report stated that nearly 1.27 million deaths worldwide occur due to bacterial AMR in 2019 [45]. AMR occurs when diverse microorganisms such as bacteria, viruses, fungi, and parasites evolve and develop resistance mechanisms against various standard treatments. One of the main reasons for growing AMR is the misuse and overuse of antimicrobials in humans, plants, and animals. To tackle this complex issue, numerous projects and plans have been adopted worldwide; for example, the Global Action Plan (GAP) on AMR was adopted by multiple nations during the 2015 World Health Assembly in order to come up with one global plan.

Another promising development to tackle AMR is the use of AMPs, which are molecules that occur naturally and target pathogens by disrupting their membranes. In the last few decades, researchers have been trying to characterize and develop new AMPs (natural and modified) with special emphasis on optimizing their structure, potency, and specificity. The FDA recently approved seven AMPs to treat different types of bacterial infections [46]. They are Gramicidin, Daptomycin, Colistin, Vancomycin, Oritavancin, Dalbavancin, and Telvancin. With advancements in next-generation sequencing techniques and artificial intelligence, scientists are exploring the genome and metagenomes to characterize new AMPs with potential therapeutics. Santos-Junior et al. recently reported that they characterized ∼8.6 million AMPs by looking at 87,920 prokaryotic genomes and 63,410 metagenomes using in silico techniques. Santos-Junior et al. synthesized 100 AMPs and demonstrated their effectiveness against ESKAPEE (*Enterococcus faecium, Staphylococcus aureus, Klebsiella pneumoniae, Acinetobacter baumannii, Pseudomonas aeruginosa, Enterobacter spp.,* and *Escherichia coli*) pathogens, a recognized public health concern [17]. Likewise, Wan et al. implemented a deep learning approach to discover antibiotic peptides by mining proteomes of all the available extinct organisms [47].

In the current study, we performed a meta-analysis to predict class-specific activity of the peptides using 8.6 million AMPs provided in the AMP Sphere database. Class-specific peptides are very important because they target specific pathogens (bacteria, viruses, fungi, etc.) better than general AMPs. This means they are more effective, less likely to have side effects, and less likely to cause resistance by focusing on different microbial mechanisms. We filtered the 8.6 million peptides based on certain criteria (experimental validation, length, etc.) and predicted their activity as ABPs, AFPs, or AVPs using various machine learning-based online tools. Peptides predicted positively by all of the tools with high prediction scores were selected for further analyses. For negative datasets, we implemented 3 different approaches (see Methods). After that, we performed numerous analyses, such as looking at the amino acid composition, physiochemical properties, motif analysis, residue positional preference at the N and C termini, and creating class-specific machine learning models using multiple classifiers. Composition analysis showed that ABPs were rich in basic and non-polar amino acids (A, C, K, P & R); AFPs were enriched for major residues aromatic (F, W, and Y), polar (S, T, and N), and aliphatic (G) in nature, whereas AVPs were enriched in amino acids that are non-polar and aliphatic (I, L, and V) in nature. According to physiochemical analysis, AFPs have the highest molecular weight, followed by ABPs and then AVPs; ABPs were found to have the highest charge and isoelectric point, followed by AFPs and then AVPs. Positional preference analysis using two sample logos suggested distinct preferences for residues at the N and C termini for three classes of AMPs. For the N-terminus, residues such as K, A, R, and V were highly preferred at the first few positions in ABPs; L, I, V, G, and F in AFPs and AVPs. For C-terminus, we observed preference of residues K, R, V, and G in ABPs; K, W, and G in AFPs; and K, R, V, and M in AVPs. Likewise, motif analyses showed enrichment of distinct motifs in ABPs, AVPs, and AFPs.

We also developed seven distinct machine learning classifiers in a class-specific manner. The extra tree-based classifier performed best based on the amino acid composition feature, with an AUROC of 0.983 on the training dataset and 0.984 on the independent dataset for ABP prediction; an AUROC of 0.987 on the training dataset and 0.986 on the independent dataset for AFP prediction; and an AUROC of 0.988 on the training dataset and 0.989 on the independent dataset for AVP prediction. This demonstrated that this newly created dataset could be useful in predicting the activity of a peptide with high accuracy, as well as paving the way for developing more sophisticated in silico tools.

We screened the peptides further to obtain peptides with high therapeutic potential. We checked for the toxicity, allergenicity, and cell-penetrating properties of the peptides using various online servers and selected the peptides fulfilling all three properties. We computed the secondary structure content for this set of peptides using the GOR4 webserver, and lastly, we predicted the peptides’ half-lives. Top10 peptides with maximum half-lives were selected in each class, i.e., ABPs, AFPs, and AVPs, and we proposed these peptides to be the ones with the highest therapeutic potential. To further confirm the therapeutic properties of these peptides, we predicted the structure of the top 10 peptides in each class using ColabFold embedded in ChimeraX1.8 software. Post-structure prediction, we carried out molecular docking studies using the HDock webserver, and as a case example, we selected one pathogenic protein from each class (methods). We observed strong non-covalent interactions among the protein and peptide residues with high free energy and docking scores.

Overall, we implemented various *in silico* analyses to evaluate the high-throughput peptide data and were able to predict class-specific peptides with significant therapeutic properties in a class-specific manner.

## Data and Code Availability

This paper analyzes existing, publicly available data. The accession numbers for the datasets used are listed in the key resources table.

Any additional information required to reanalyze the data reported in this paper is available from the lead contact upon request.

## Author contributions

PA collected and processed the dataset. PA, AP, and RRL performed the analysis. AP performed the structure prediction and molecular docking analysis. SP performed the machine learning analysis. RRL created the tables and figures. PA, RRL and AP wrote the manuscript. PA conceived the idea and supervised the study.

## Acknowledgement

This research was supported by SRM Institute of Science & Technology, Kattankulathur, India.

## Declaration of generative AI and AI-assisted technologies in the writing process

During the preparation of this work the author do not used any AI or AI-assisted technologies.

