## Supplementary Figures for "High Throughput Meta-analysis of Antimicrobial Peptides for Characterizing Class Specific Therapeutic Candidates: An *in-silico* Approach"

Piyush Agrawal, Ph.D.

Division of Medical Research, SRM Medical College Hospital & Research Centre, SRMIST, Kattankulathur, Chennai, India-603203

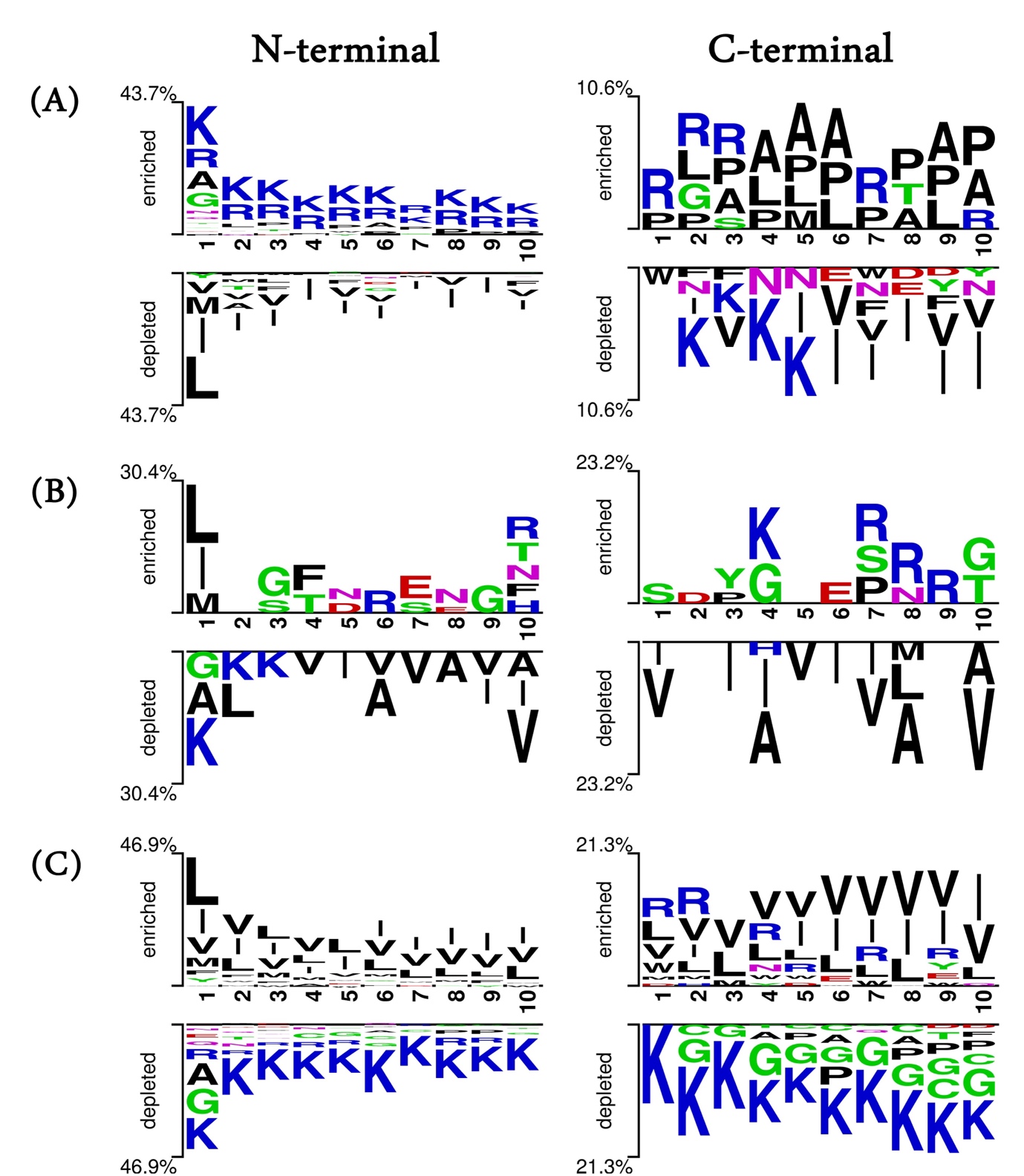

**Supplementary Figure S1. Positional Preference Analysis.** Two Sample Logo analysis showing the amino acid residue preference for the first 10 residues from N-terminal and C-terminal in ABPs and non-ABPs (A); AFPs and non-AFPs (B); and AVPs and non-AVPs (C); for the dataset HQ_DS1.

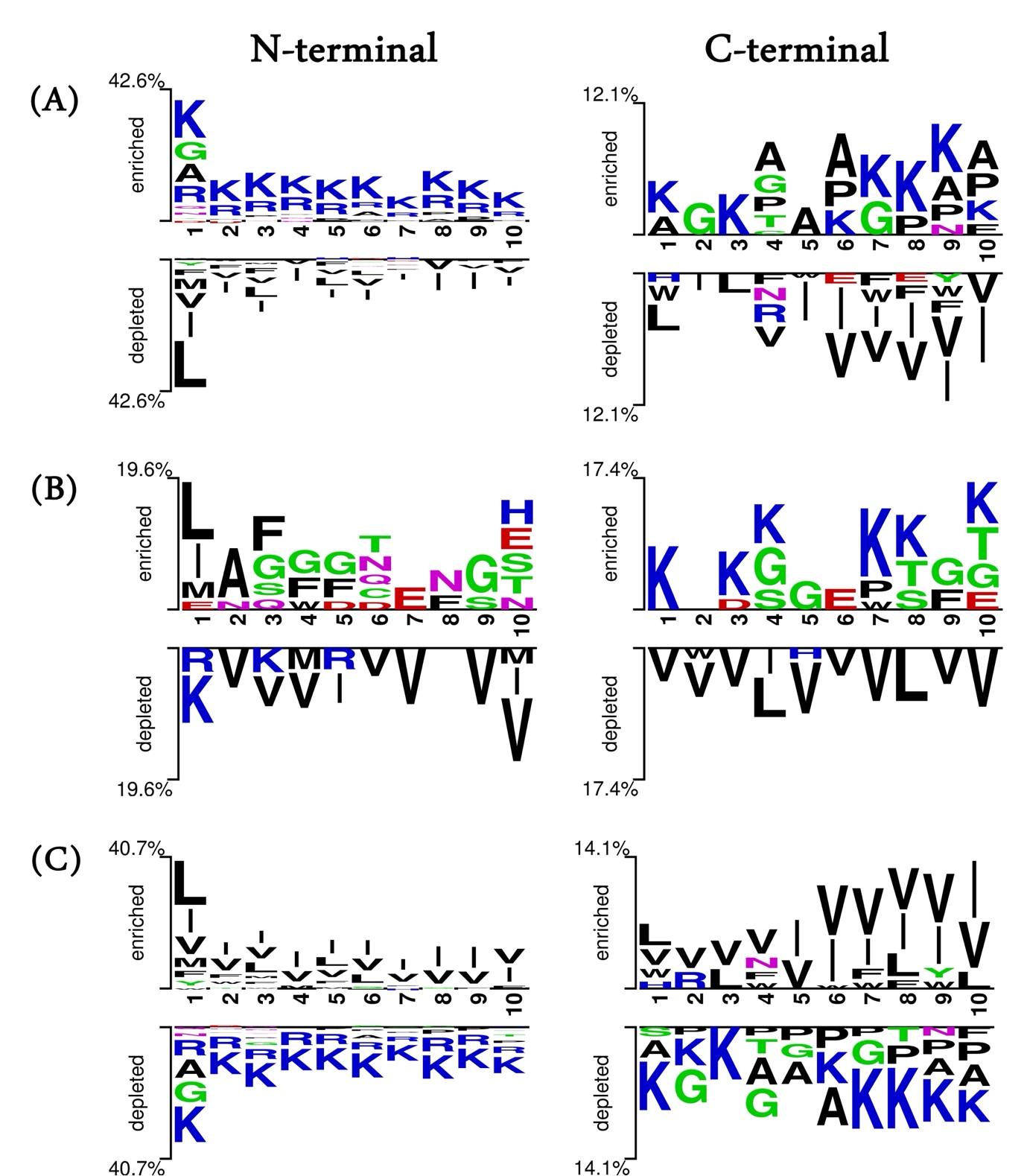

**Supplementary Figure S2. Positional Preference Analysis.** Two Sample Logo analysis showing the amino acid residue preference for the first 10 residues from N-terminal and C-terminal in ABPs and non-ABPs (A); AFPs and non-AFPs (B); and AVPs and non-AVPs (C); for the dataset HQ_DS3.

| 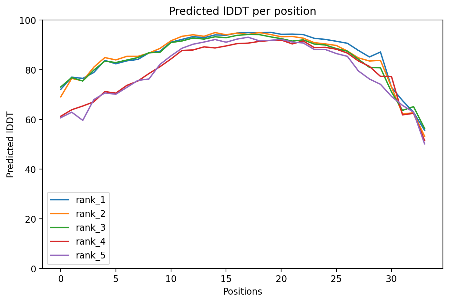 | 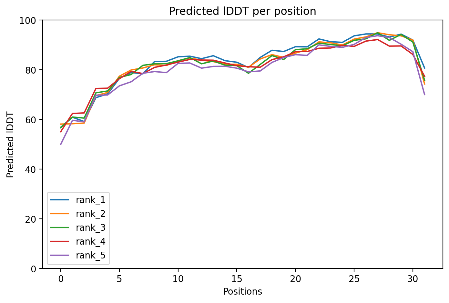 | 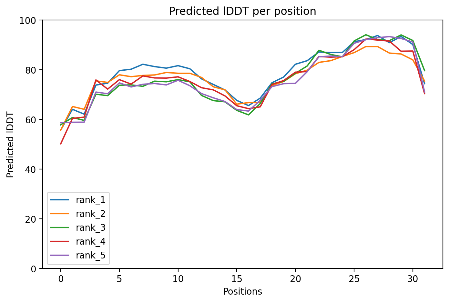 |
| --- | --- | --- |
| **(a) Bpep-01** | **(b) Bpep-02** | **(c) Bpep-03** |
| 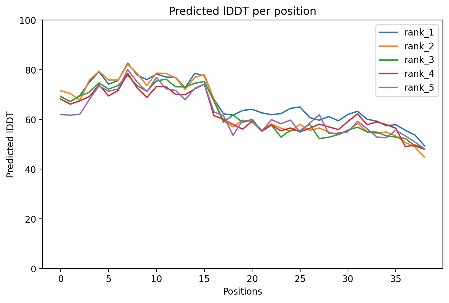 | 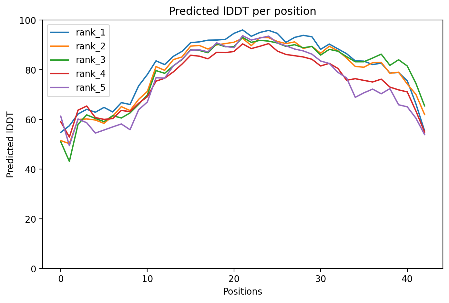 | 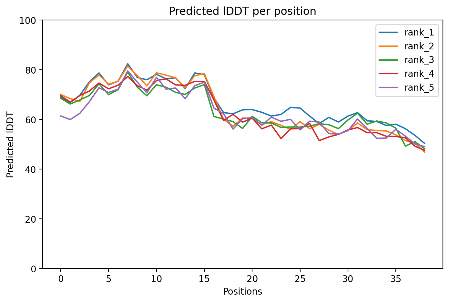 |
| **(d) Bpep-04** | **(e) Bpep-05** | **(f) Bpep-06** |
| 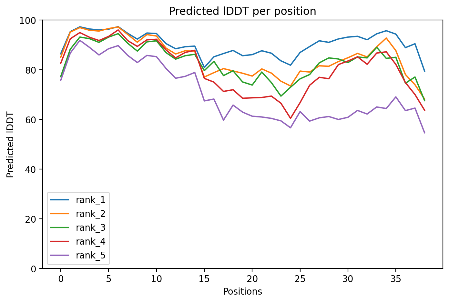 | 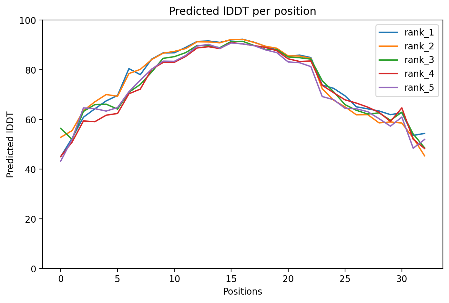 | 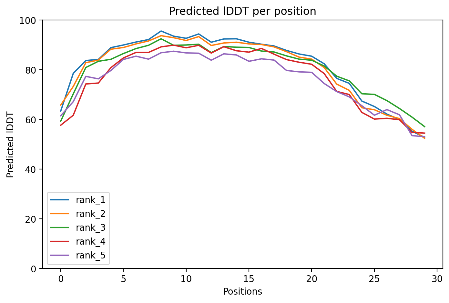 |
| **(g) Bpep-07** | **(h) Bpep-08** | **(i) Bpep-09** |
| 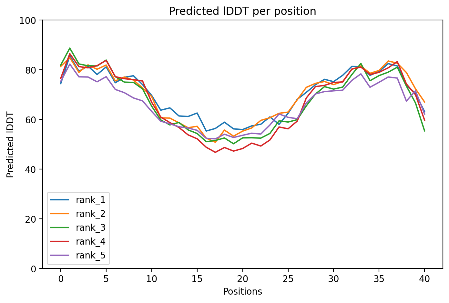 |  |  |
| **(j) Bpep-10** |  |  |

**Supplementary Figure S3(a).** The plot represents the pLDDT per residue position of the input anti-bacterial peptide sequences, for each of the five predicted models. pLDDT > 90 are expected to be modelled to high accuracy; 90 > pLDDT > 70 are expected to be modelled well; 70 > pLDDT > 50 are expected to be modelled not so well.

| 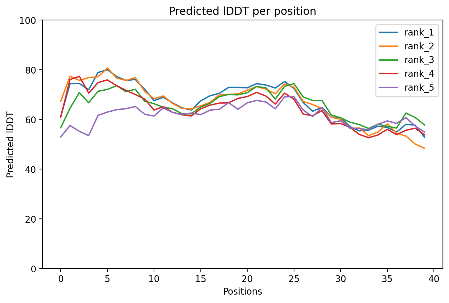 | 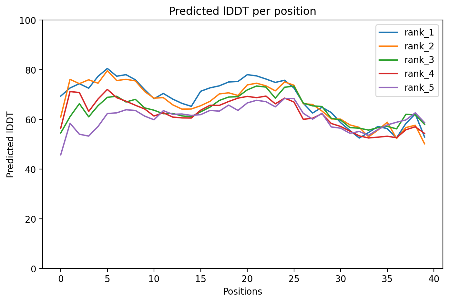 | 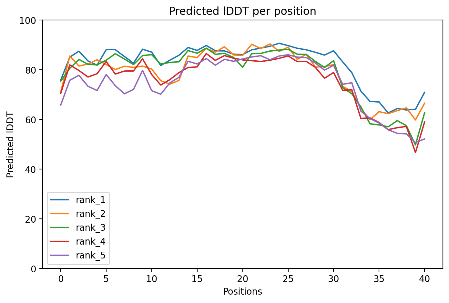 |
| --- | --- | --- |
| **(a) Fpep-01** | **(b) Fpep-02** | **(c) Fpep-03** |
| 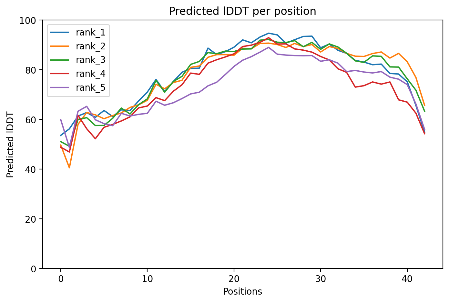 | 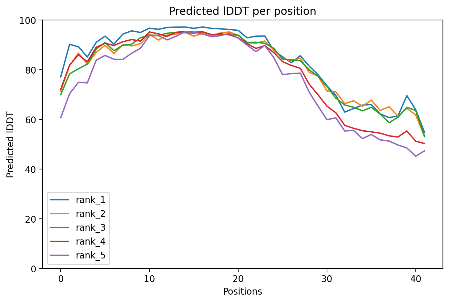 | 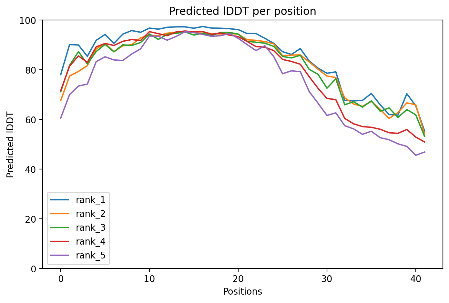 |
| **(d) Fpep-04** | **(e) Fpep-05** | **(f) Fpep-06** |
| 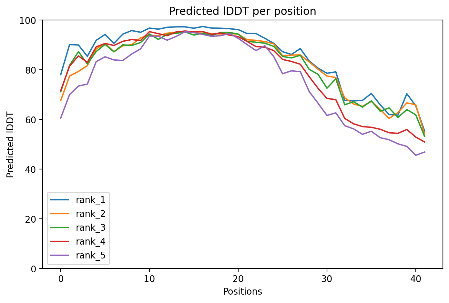 | 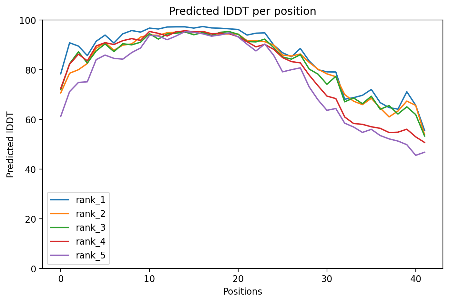 | 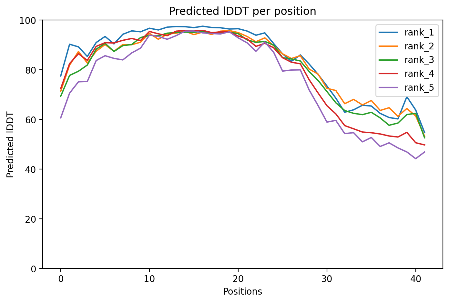 |
| **(g) Fpep-07** | **(h) Fpep-08** | **(i) Fpep-09** |
| 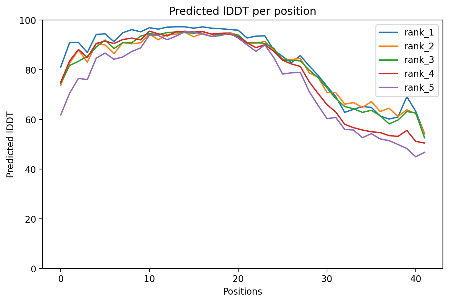 |  |  |
| **(j) Fpep-10** |  |  |

**Supplementary Figure S3(b).** The plot represents the pLDDT per residue position of the input anti-fungal peptide sequences, for each of the five predicted models. pLDDT > 90 are expected to be modelled to high accuracy; 90 > pLDDT > 70 are expected to be modelled well; 70 > pLDDT > 50 are expected to be modelled not so well.

| 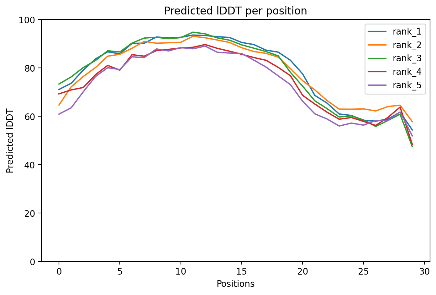 | 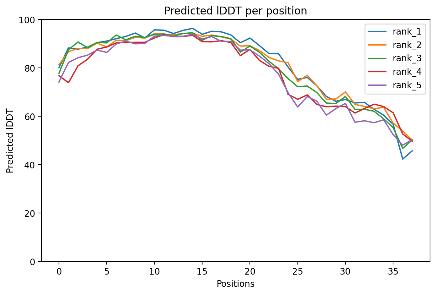 | 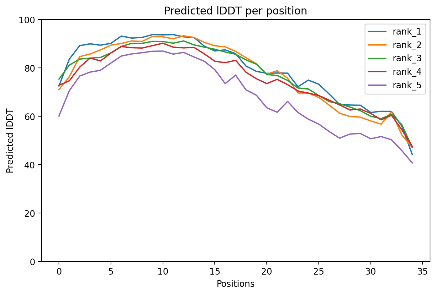 |
| --- | --- | --- |
| **(a) Vpep-01** | **(b) Vpep-02** | **(c) Vpep-03** |
| 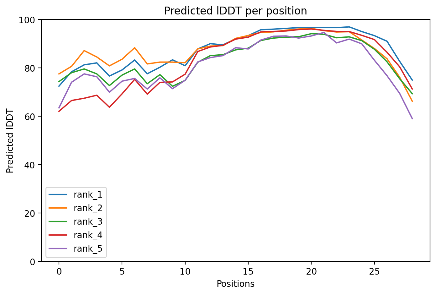 | 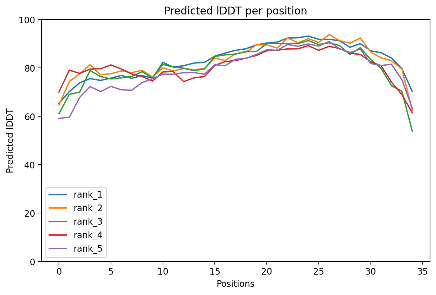 | 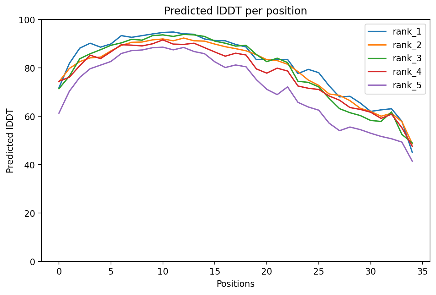 |
| **(d) Vpep-04** | **(e) Vpep-05** | **(f) Vpep-06** |
| 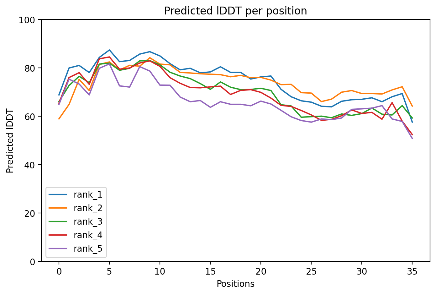 | 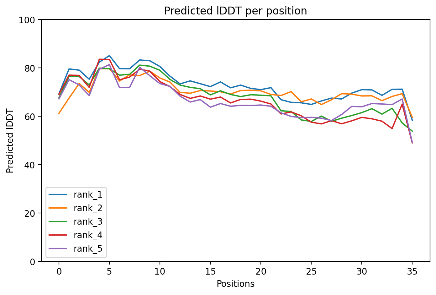 | 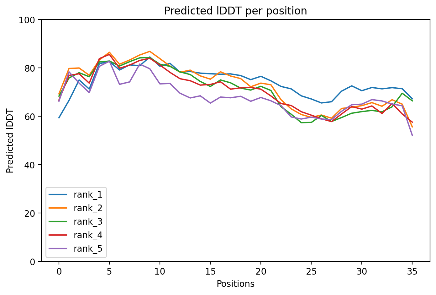 |
| **(g) Vpep-07** | **(h) Vpep-08** | **(i) Vpep-09** |
| 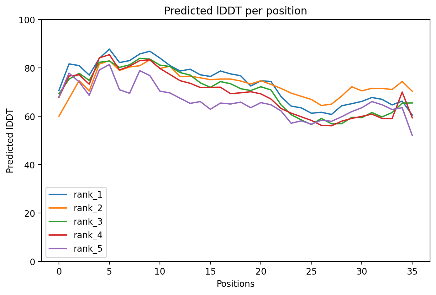 |  |  |
| **(j) Vpep-10** |  |  |

**Supplementary Figure S3(c).** The plot represents the pLDDT per residue position of the input anti-viral peptide sequences, for each of the five predicted models. pLDDT > 90 are expected to be modelled to high accuracy; 90 > pLDDT > 70 are expected to be modelled well; 70 > pLDDT > 50 are expected to be modelled not so well.

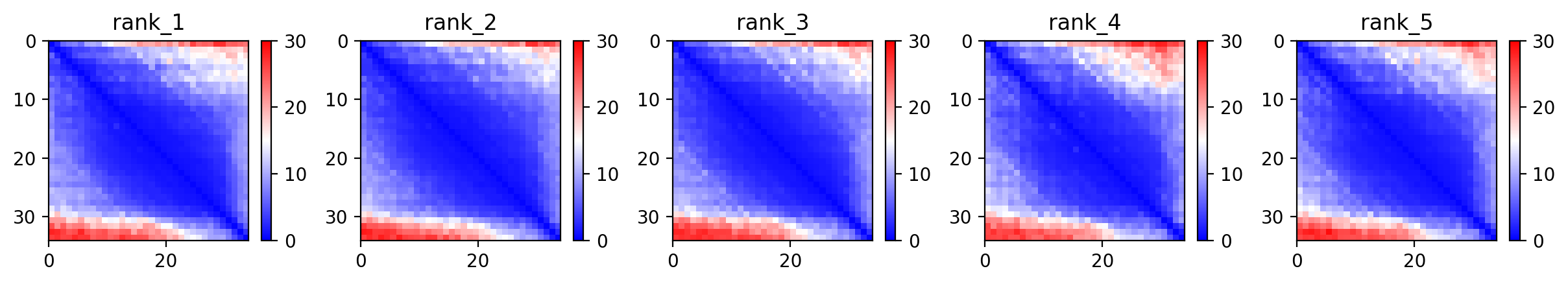

(a) **Bpep-01**

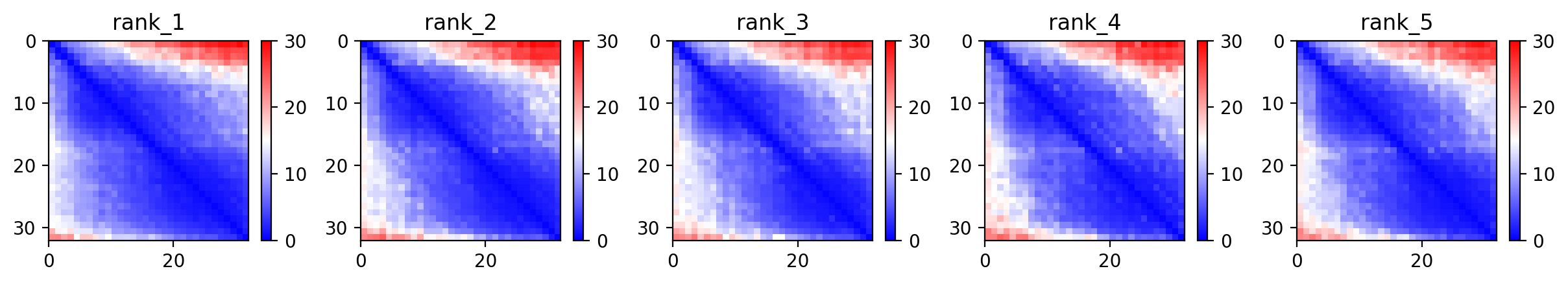

**(b) Bpep-02**

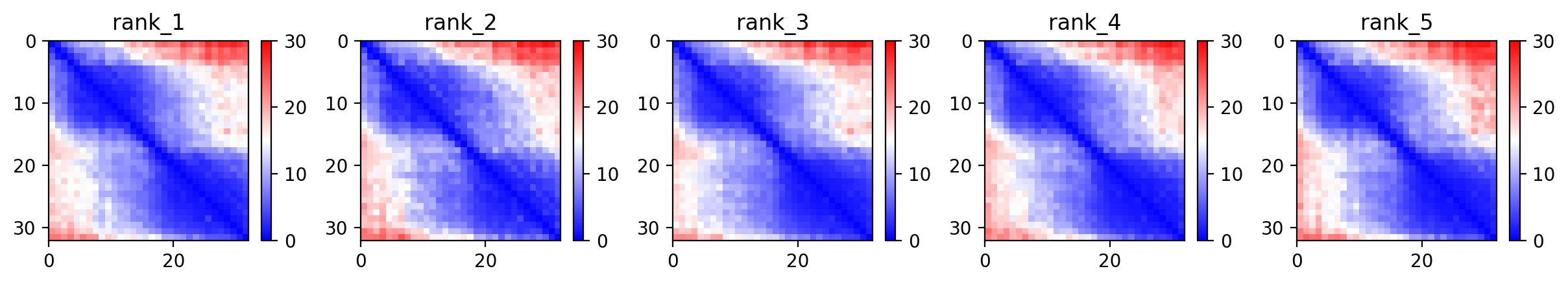

**(c) Bpep-03**

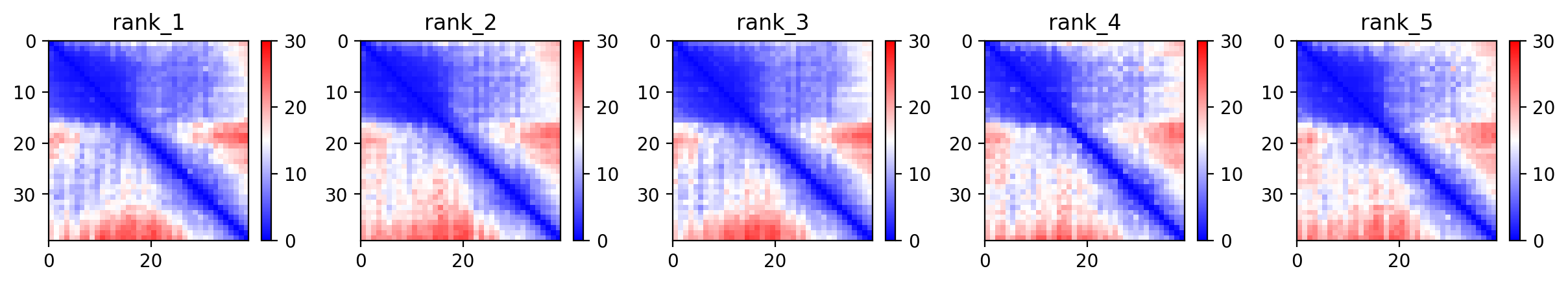

**(d) Bpep-04**

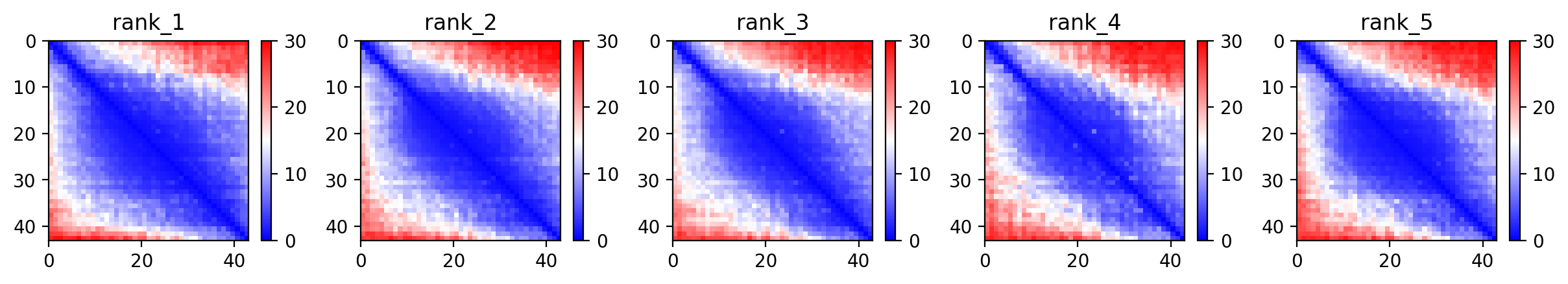

**(e) Bpep-05**

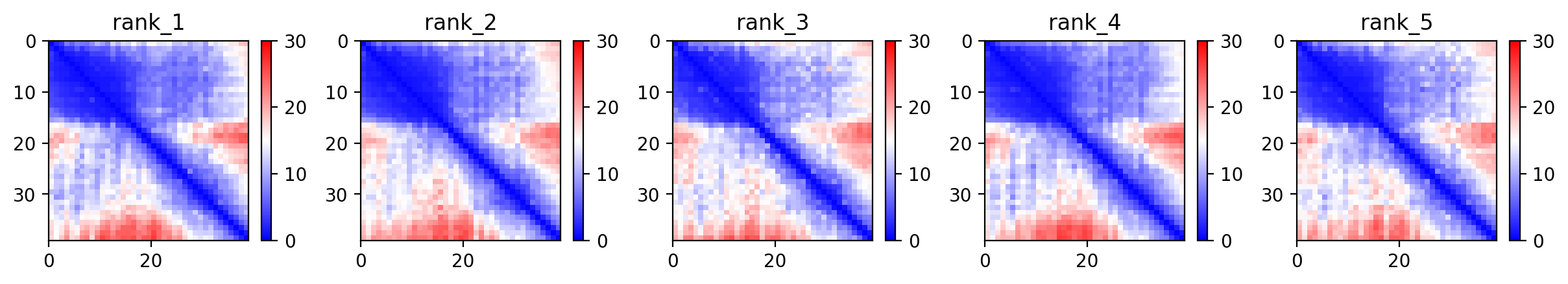

**(f) Bpep-06**

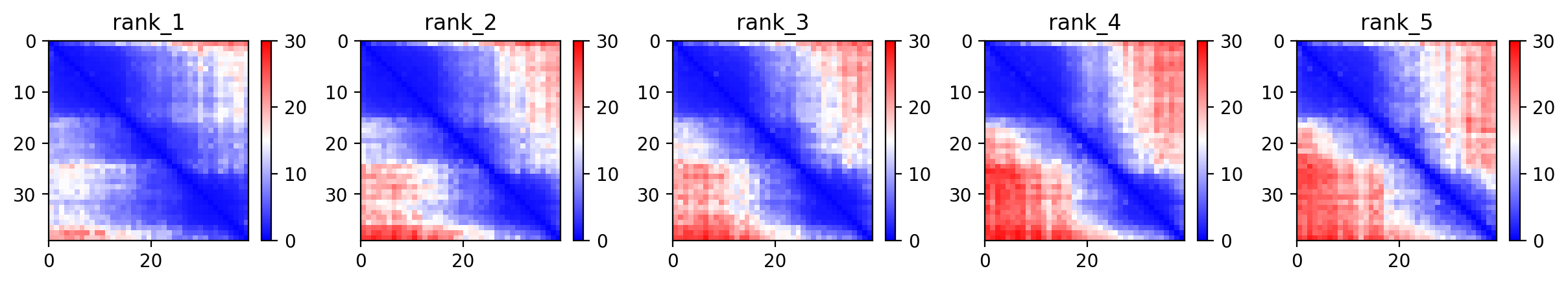

**(g) Bpep-07**

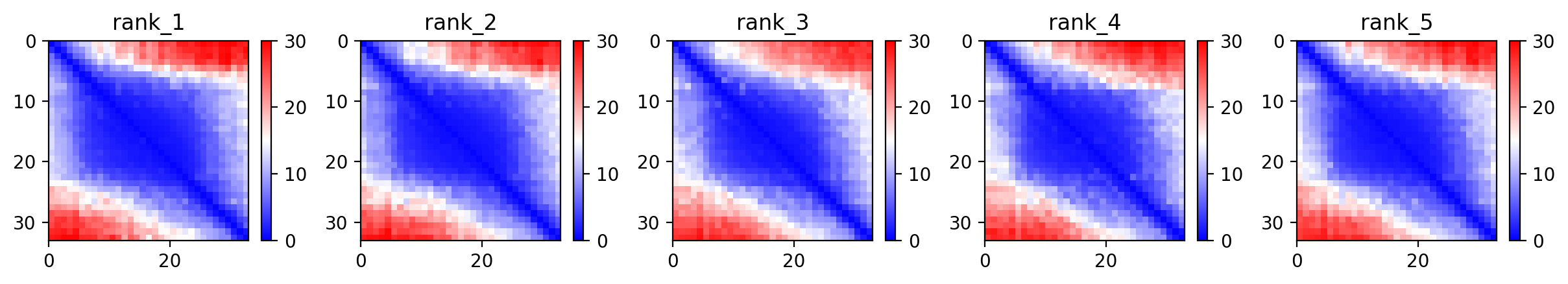

**(h) Bpep-08**

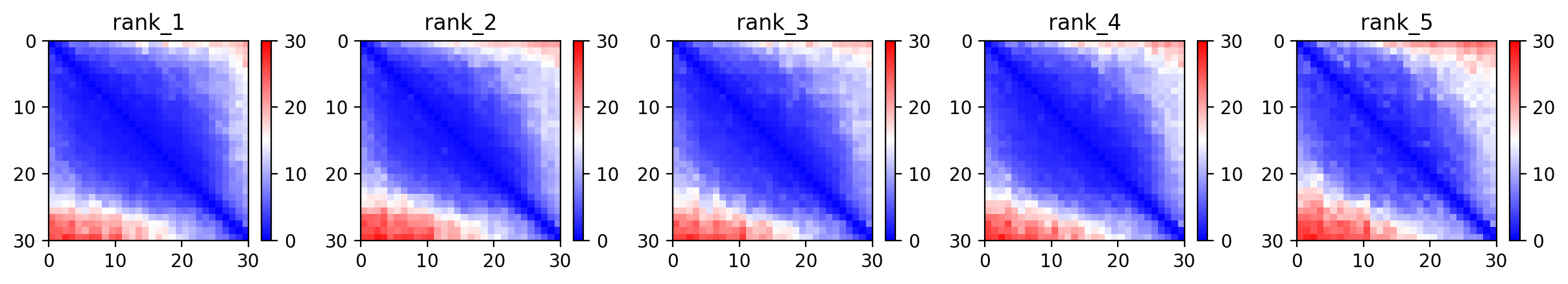

**(i) Bpep-09**

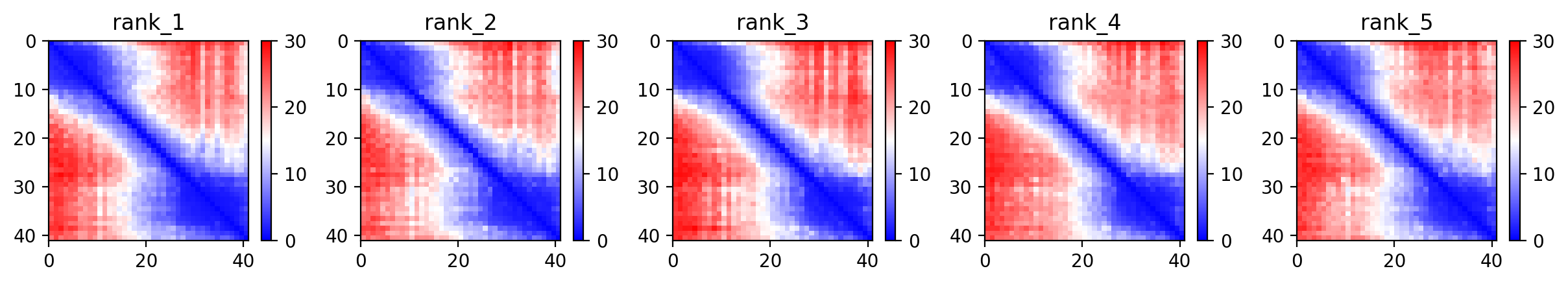

**(j) Bpep-10**

**Supplementary Figure S4(a).** The heat maps represent the predicted alignment error between each residue in the model. The colour scale contains three colours to further accentuate the contrast between the high confidence regions and the low confidence regions. Higher blue and lower red areas correspond to higher degree of accuracy.

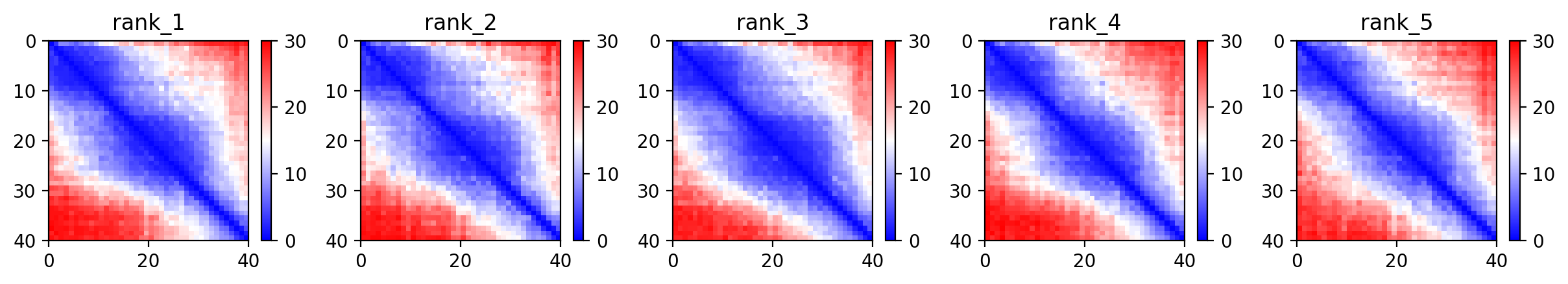

**(a) Fpep-01**

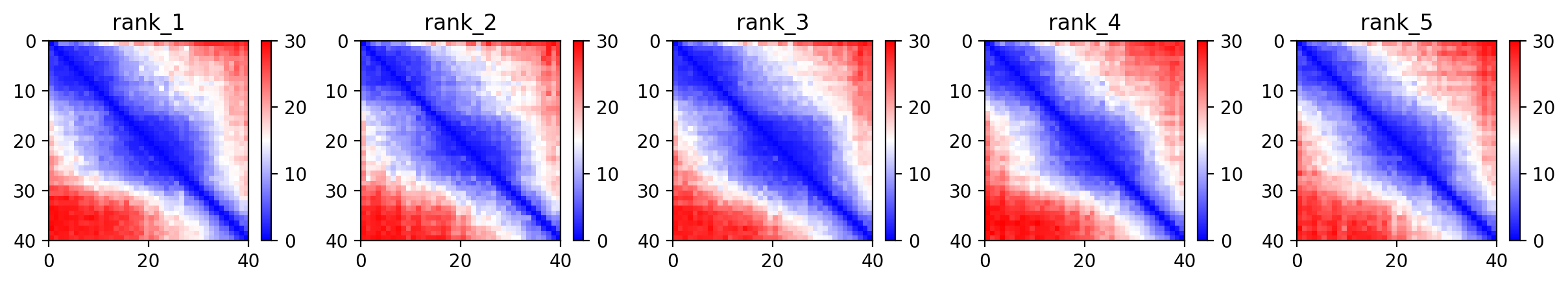

**(b) Fpep-02**

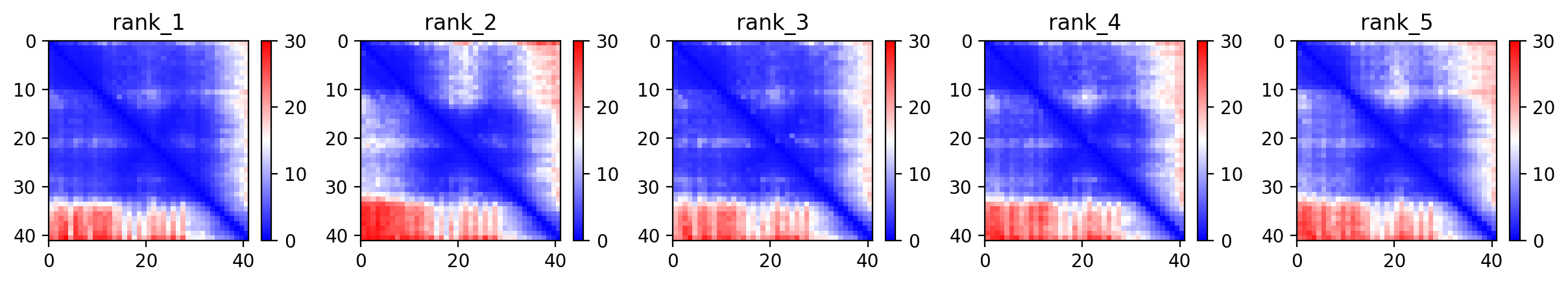

**(c) Fpep-03**

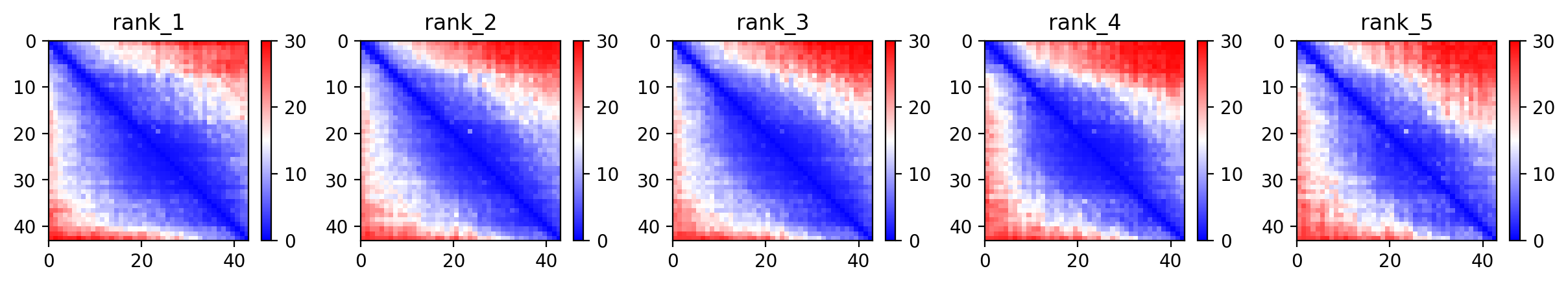

**(d) Fpep-04**

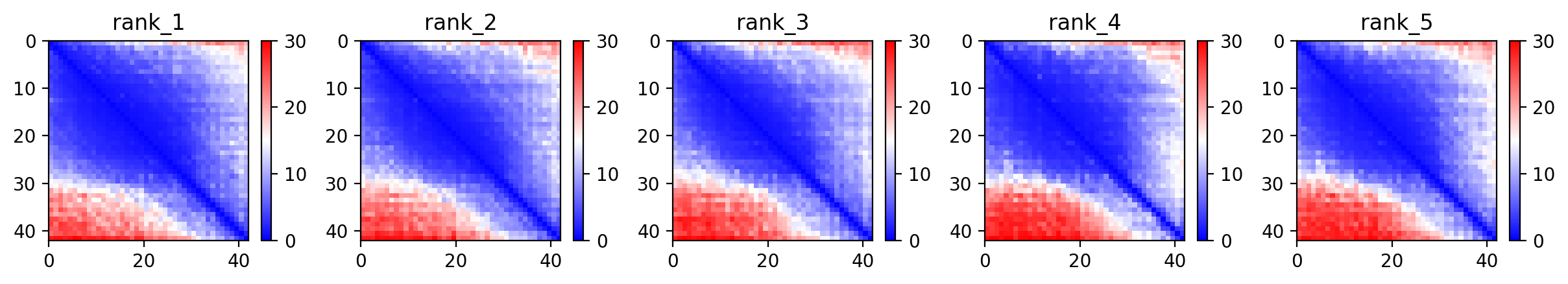

**(e) Fpep-05**

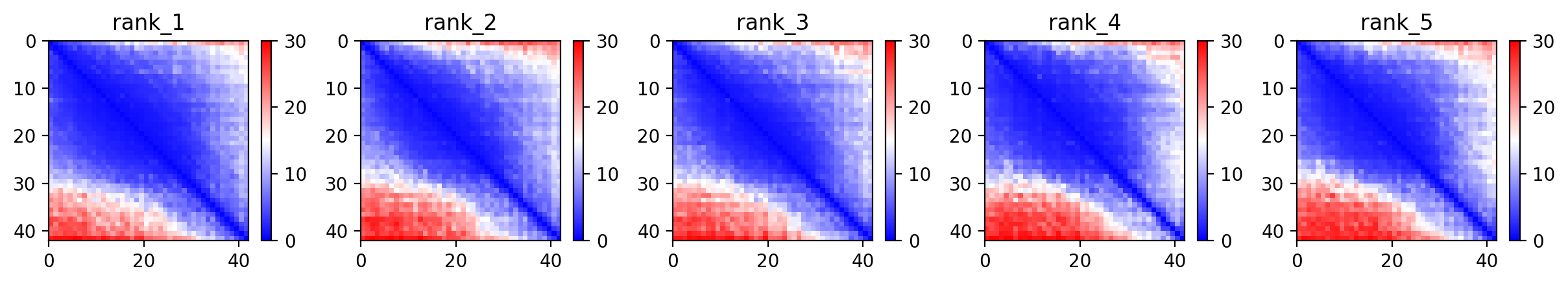

**(f) Fpep-06**

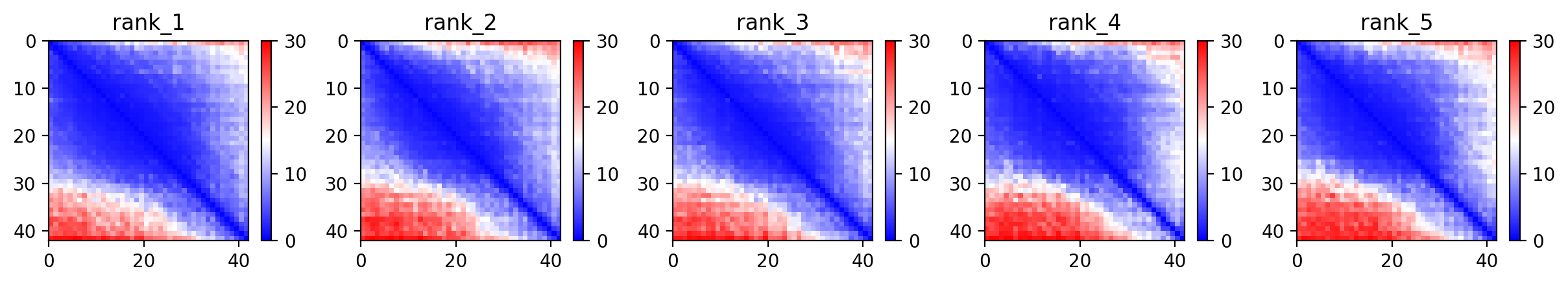

**(g) Fpep-07**

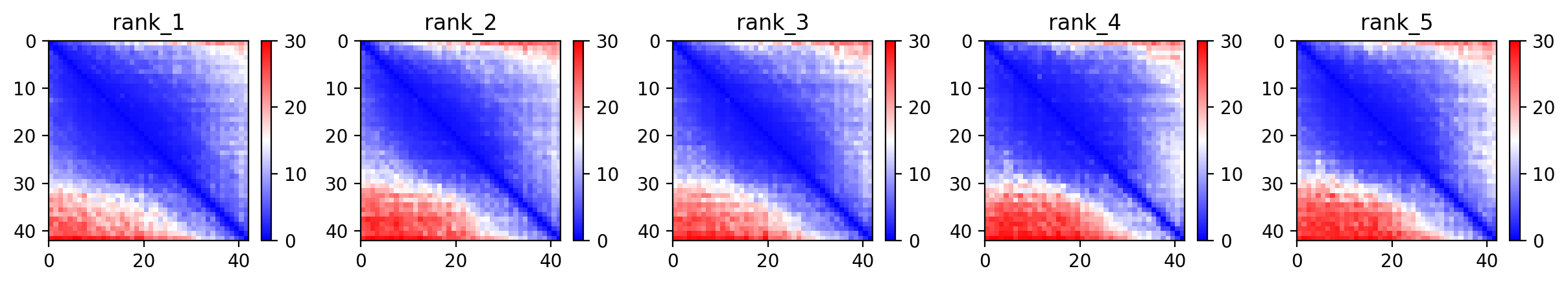

**(h) Fpep-08**

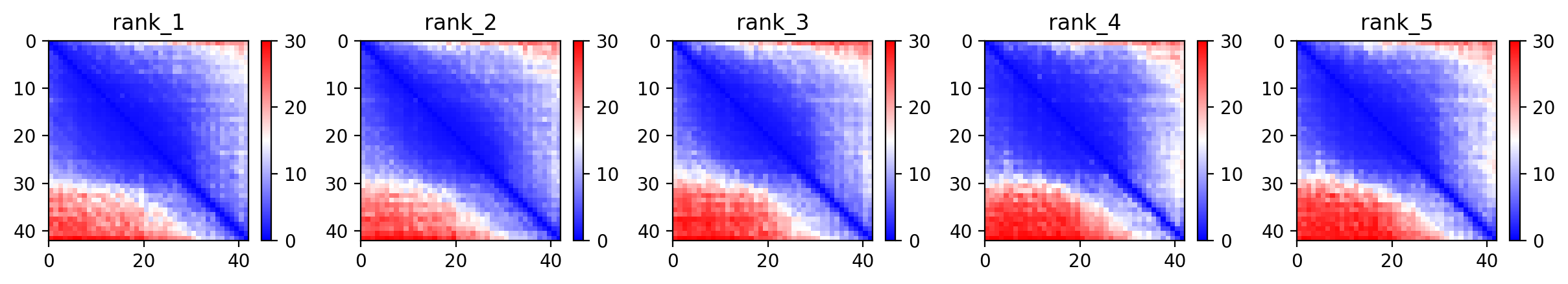

**(i) Fpep-09**

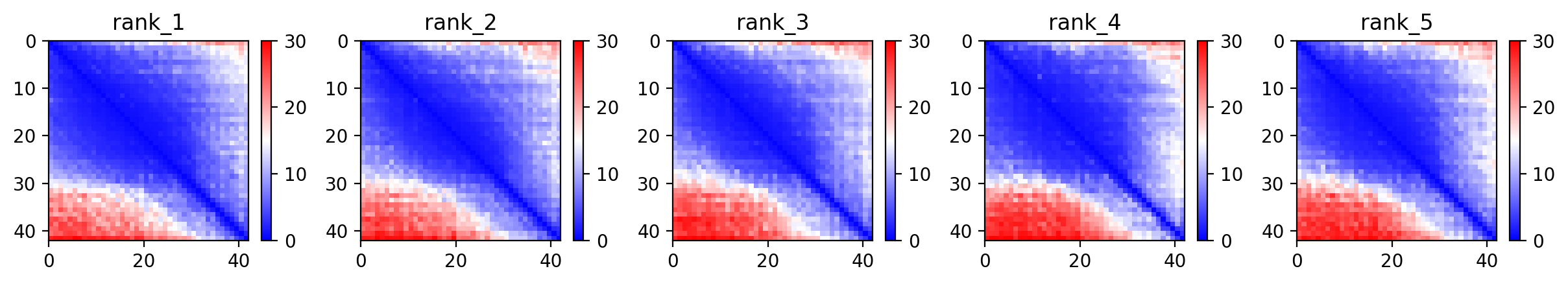

**(j) Fpep-10**

**Figure S4(b).** The heat maps represent the predicted alignment error between each residue in the model. The colour scale contains three colours to further accentuate the contrast between the high confidence regions and the low confidence regions. Higher blue and lower red areas correspond to higher degree of accuracy.

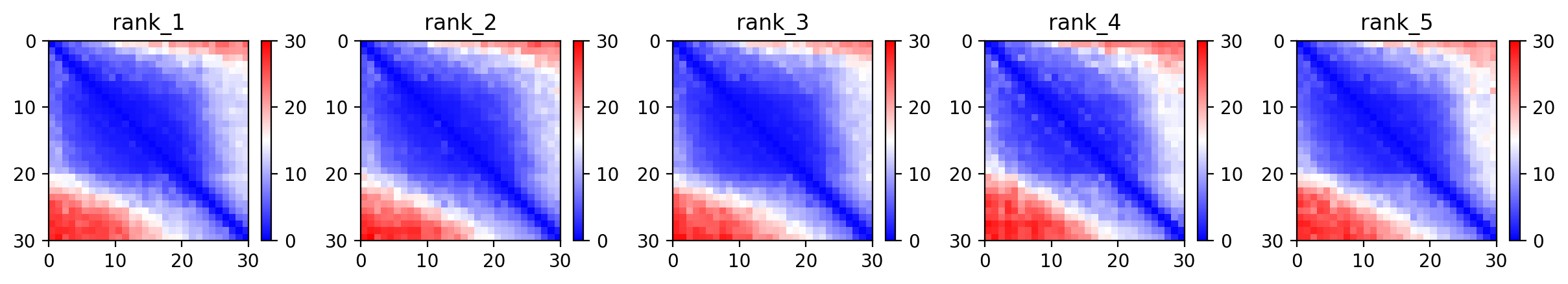

**(a) Vpep-01**

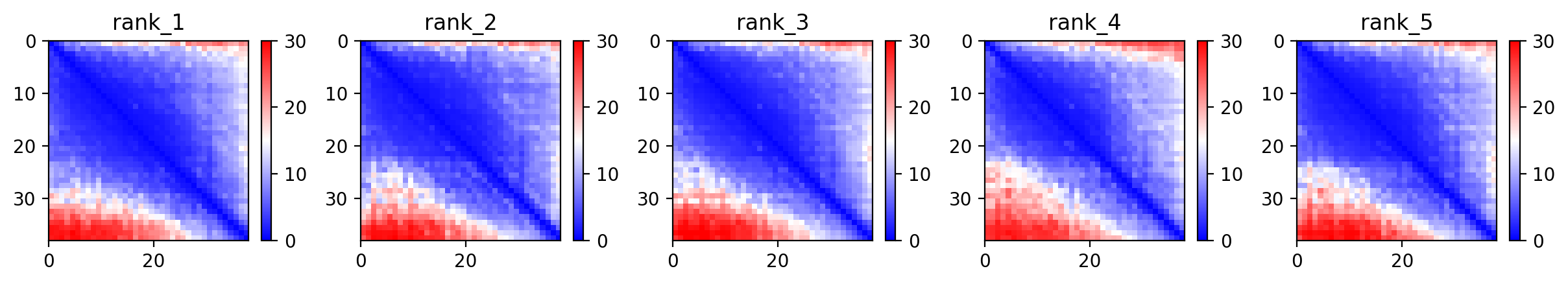

**(b) Vpep-02**

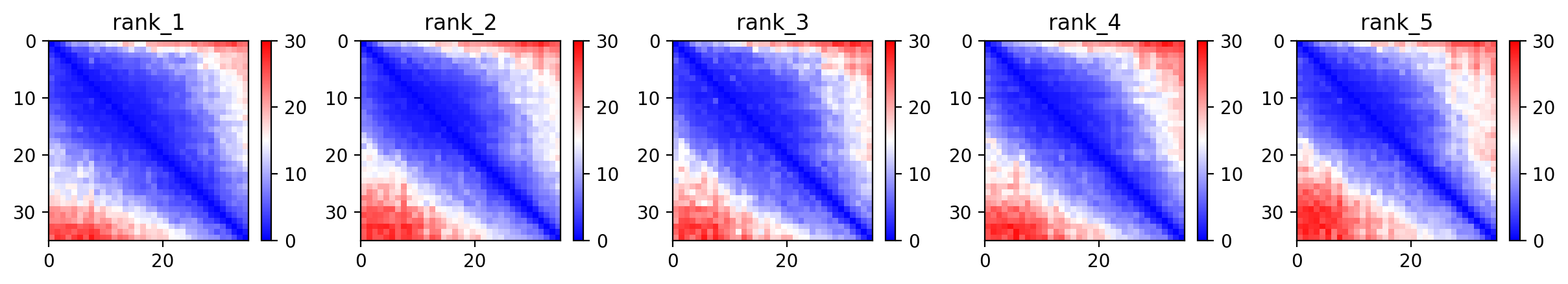

**(c) Vpep-03**

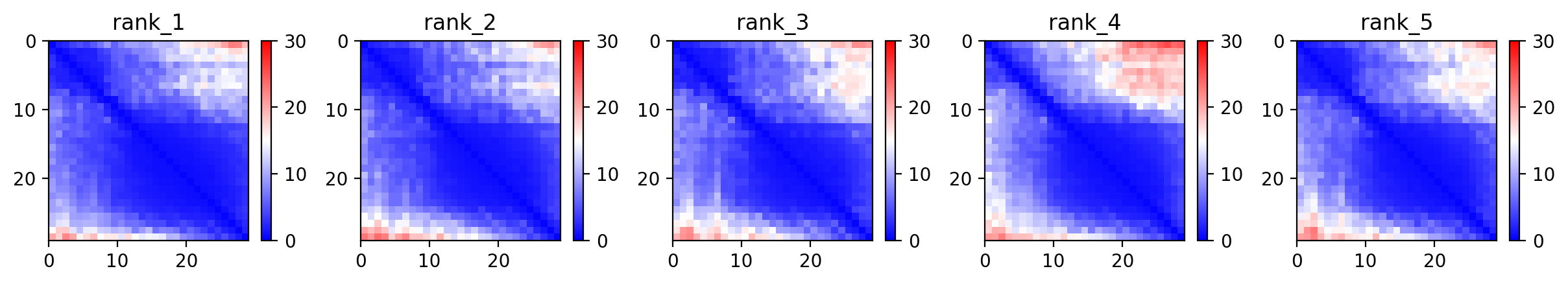

**(d) Vpep-04**

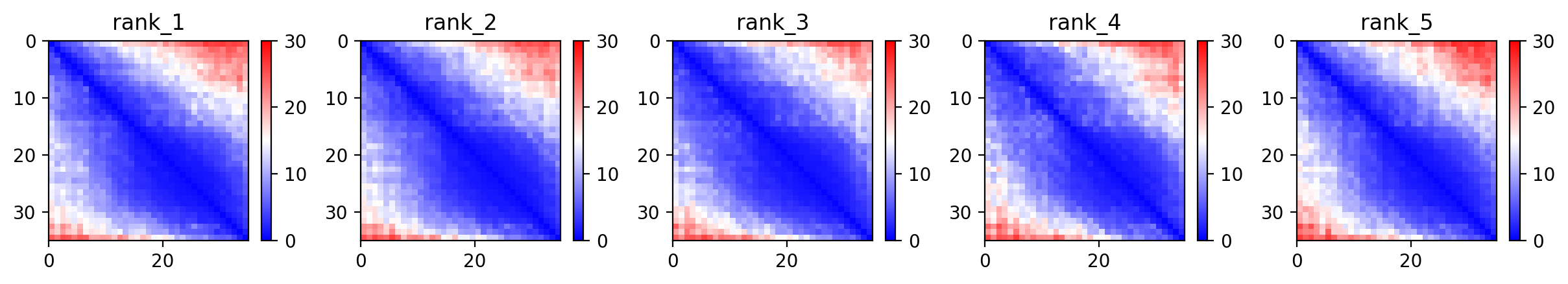

**(e) Vpep-05**

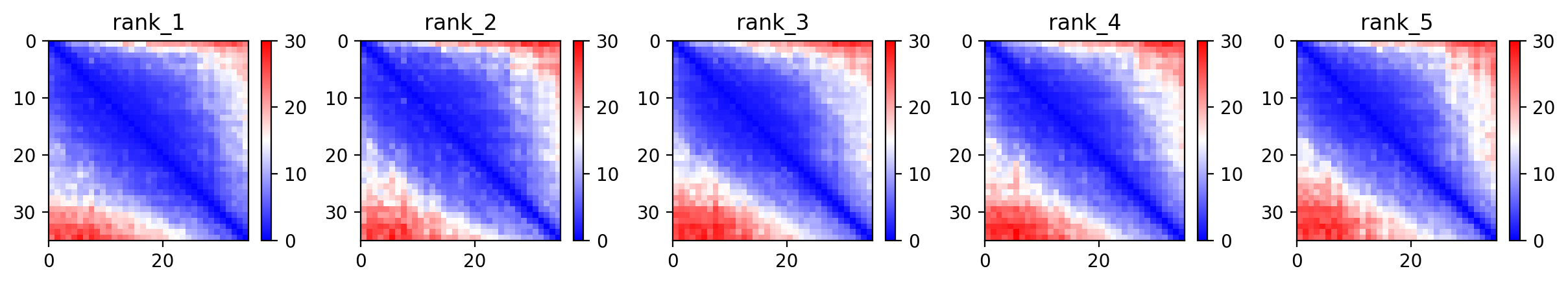

**(f) Vpep-06**

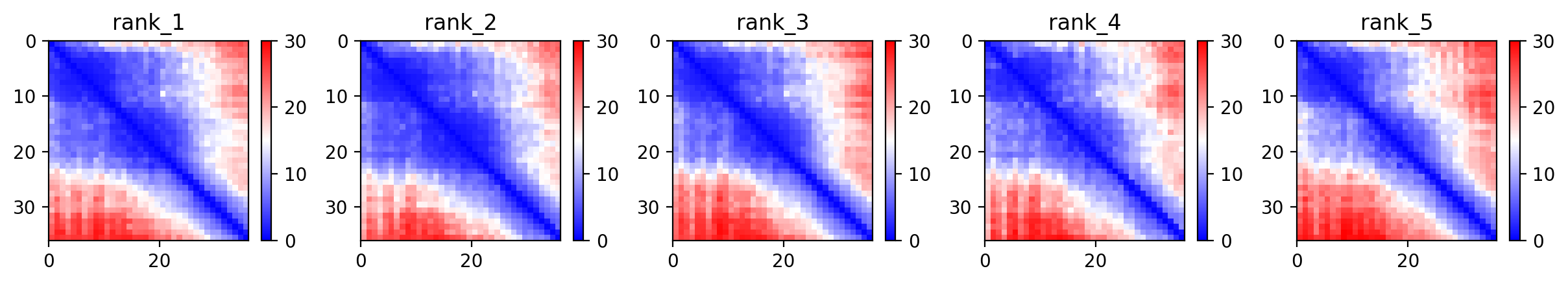

**(g) Vpep-07**

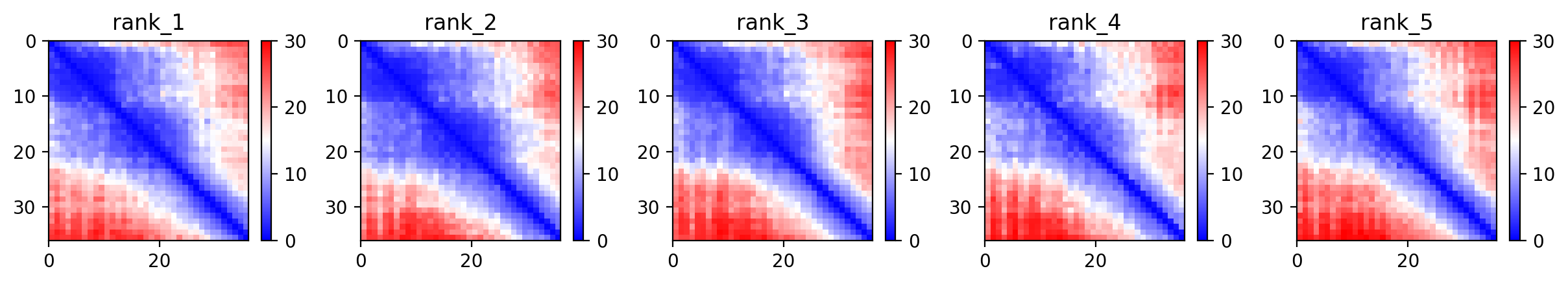

**(h) Vpep-08**

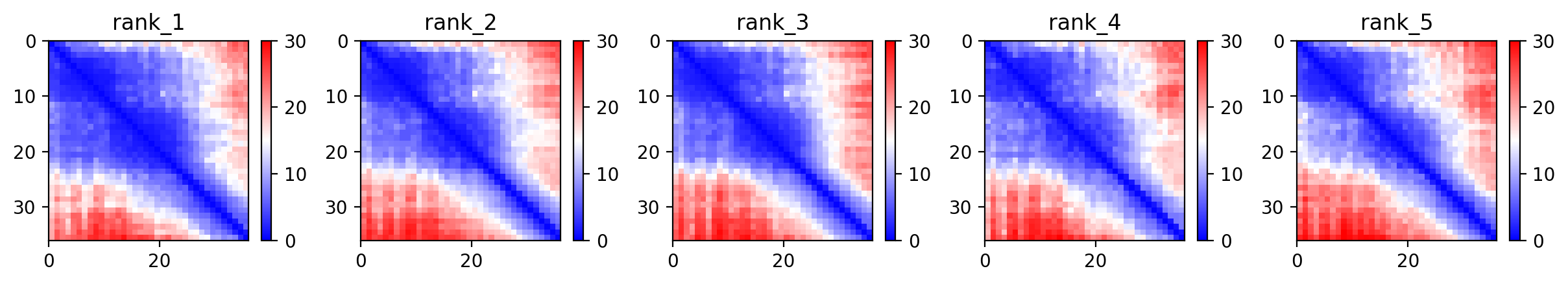

**(i) Vpep-09**

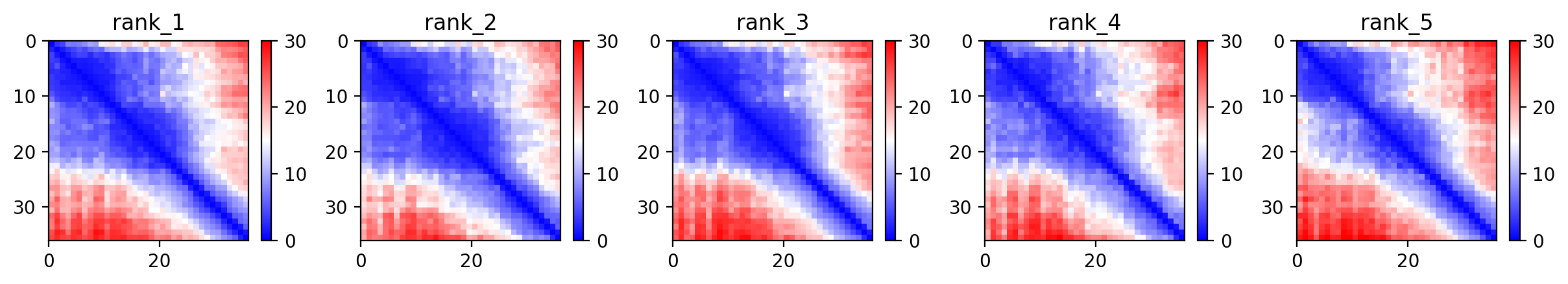

**(j) Vpep-10**

**Supplementary Figure S4(c).** The heat maps represent the predicted alignment error between each residue in the model. The colour scale contains three colours to further accentuate the contrast between the high confidence regions and the low confidence regions. Higher blue and lower red areas correspond to higher degree of accuracy.
